# Anesthetics fragment hippocampal network activity, alter spine dynamics and affect memory consolidation

**DOI:** 10.1101/2020.06.05.135905

**Authors:** Wei Yang, Mattia Chini, Jastyn A. Pöpplau, Andrey Formozov, Alexander Dieter, Patrick Piechocinski, Cynthia Rais, Fabio Morellini, Olaf Sporns, Ileana L. Hanganu-Opatz, J. Simon Wiegert

## Abstract

General anesthesia is characterized by reversible loss of consciousness accompanied by transient amnesia. Yet, long-term memory impairment is an undesirable side-effect. How different types of general anesthetics (GAs) affect the hippocampus, a brain region central to memory formation and consolidation, is poorly understood. Using extracellular recordings, chronic 2-photon imaging and behavioral analysis, we monitor the effects of isoflurane (Iso), medetomidine/midazolam/fentanyl (MMF), and ketamine/xylazine (Keta/Xyl) on network activity and structural spine dynamics in the hippocampal CA1 area of adult mice. GAs robustly reduced spiking activity, decorrelated cellular ensembles, albeit with distinct activity signatures, and altered spine dynamics. CA1 network activity under all three anesthetics was different to natural sleep. Iso anesthesia most closely resembled unperturbed activity during wakefulness and sleep, and network alterations recovered more readily than with Keta/Xyl and MMF. Correspondingly, memory consolidation was impaired after exposure to Keta/Xyl and MMF, but not Iso. Thus, different anesthetics distinctly alter hippocampal network dynamics, synaptic connectivity, and memory consolidation, with implications for GA strategy appraisal in animal research and clinical settings.

## INTRODUCTION

General anesthesia is a drug-induced, reversible behavioral condition encompassing unconsciousness, amnesia, sedation, immobility, and analgesia [1, 2]. Together, these aspects represent a state where surgery can be tolerated without the requirement for further drugs [2]. The behavioral effects of GAs are dose-dependent. At clinical (i.e. highest) dosage, they should induce unconsciousness, even though experimental evidence of this phenomenon is challenging to collect (in the absence of a verifiable consciousness theory). At lower doses, some GAs cause unresponsiveness and loss of working memory, phenomena that have both been hypothesized to potentially confound the apparent loss of consciousness [3, 4]. At much lower doses still, GAs cause profound retrograde amnesia. When general anesthesia fails to induce such behavioral effects, intraoperative awareness ensues, a condition that is associated with long-term adverse health consequences [5]. While loss of memory is required during anesthesia administration, so that no memories of the surgical procedure are formed [1, 6], long-term impairment of retrograde or anterograde memories is not desired. Although general anesthesia is generally considered a safe procedure, growing literature points to the possibility of long-term negative effects on the central nervous system [7]. This is particularly true for specific categories of patients, such as the elderly, infants and children [7]. Among the observed side effects, the most common are post-operative cognitive dysfunction syndromes, including post-operative delirium and post-operative cognitive decline. Post-operative cognitive disturbances are positively correlated with the duration of anesthesia and a single exposure to GAs can cause retrograde and anterograde memory deficits that persist for days to weeks in rodent models [8]. These aspects point to a generalized action of GAs on the memory system.

Given that amnesia is a fundamental part of general anesthesia and that the hippocampus controls memory formation and consolidation, it is important to understand how anesthetics affect hippocampal function and how this compares to sleep – a naturally occurring state of unconsciousness. Together with the subiculum, the CA1 area constitutes the main hippocampal output region. CA1 pyramidal cells receive excitatory synaptic input mainly from CA3 (in strata oriens & radiatum) and layer 3 of entorhinal cortex (in stratum lacunosum moleculare), relaying information about the internal state of the animal and sensory inputs from the external environment, respectively [9]. Inputs along these pathways are processed in an integrative manner in CA1 [10]. Thus, CA1 pyramidal cells have been suggested to be a site of sensory integration, with synaptic spines as a possible location of memory storage [11-14]. Moreover, dynamic modulation of spine stability has been linked to synaptic plasticity [15-18]. Synaptic plasticity, in turn, underlies learning and memory formation [19], suggesting that spine turnover in the hippocampus directly reflects these processes [20, 21]. Considering the low concentrations of anesthetics required to induce amnesia, these compounds are thought of being particularly effective on the hippocampus. One possible explanation of this sensitivity is the fact that a class of γ-aminobutyric acid receptors (GABARs), which is strongly modulated by some anesthetics, is predominantly expressed in the hippocampus [22, 23]. Other anesthetics, such as ketamine, inhibit N-methyl-D-aspartate receptors (NMDARs) in a use-dependent manner and therefore may be particularly effective in inhibiting synaptic plasticity, required for the formation of episodic-like memories [24]. However, a systematic investigation of the effects of anesthetics on the hippocampus, bridging synaptic, network and behavioral levels, is still lacking.

Here, using extracellular LFP and spiking recordings and chronic 2-photon calcium and spine imaging in vivo in combination with behavioral analysis, we systematically assessed how CA1 network dynamics, synaptic structure and memory performance are affected by three commonly used combinations of GAs: isoflurane (Iso), midazolam/medetomidine/fentanyl (MMF), and ketamine in combination with xylazine (Keta/Xyl). We further measured CA1 network dynamics during wakefulness and natural sleep. Unlike sleep, all three GAs strongly reduced overall neuronal spiking compared to wakefulness. Moreover, opposite to what has been found in the neocortex [25-27], they decorrelated network activity, leading to a fragmented network state. However, the induced patterns of activity were highly distinct between the three different anesthetic conditions and recovered to the pre-anesthetic status with disparate rates. Testing the effect of repeated anesthesia on spine dynamics revealed that Keta/Xyl, the condition which most strongly affected calcium activity, significantly reduced spine turnover, leading to an overall (over)stabilization of hippocampal synapses. In contrast, Iso and MMF mildly increased spine turnover. Finally, we show that the two anesthetic conditions which induce the strongest reduction and fragmentation of CA1 network activity, Keta/Xyl and MMF, negatively influenced hippocampus-dependent memory consolidation. On the other hand, Iso, which most closely resembled unperturbed sleep and wakefulness, did not impair memory consolidation, even when maintained over time periods matching the longer recovery phase of Keta/Xyl or MMF. Thus, different anesthetics, despite inducing a similar physiological state, strongly differ in their effects on synaptic stability, hippocampal network activity, and memory consolidation.

## RESULTS

### Iso, Keta/Xyl and MMF induce distinct patterns of network activity

Iso, Keta/Xyl and MMF have distinct molecular targets and modes of action in the brain. We therefore hypothesized that electrical activity in the hippocampus might be uniquely altered by the three anesthesia strategies. To test this hypothesis, we investigated local field potentials (LFPs) and firing of individual neurons (single-unit activity, SUA) extracellularly recorded in the CA1 area of the dorsal hippocampus (dCA1) during wakefulness, followed by 45 min of anesthesia and 45 min of recovery (Fig. 1A, S1A). We found that the anesthetics differently affected population activity, inducing characteristic modulation of various frequency bands (Fig. 1B). During wakefulness, LFP power in CA1 was highest in the theta (4-12 Hz) and low-gamma (40-60 Hz) frequency bands (Fig. S1B). Exposure to 2-2.5% Iso led to a strong reduction of LFP power > 4 Hz within the first 2 minutes, which was accompanied by complete loss of mobility of the animal (Fig. 1C, S1B,C). Similarly, MMF injection promptly decreased LFP power in the same frequency bands. In contrast, Keta/Xyl increased power across all frequencies during the first 10 min after injection, the most prominent effect being observed for activity at 5-30 Hz. This is consistent with previous reports, finding enhanced theta and low-gamma power in CA1 of rats under ketamine anesthesia [28]. The initial LFP power increase was followed by a gradual, significant decrease of 30-100 Hz activity (Fig. 1C, S1B,C).

**Figure 1:**
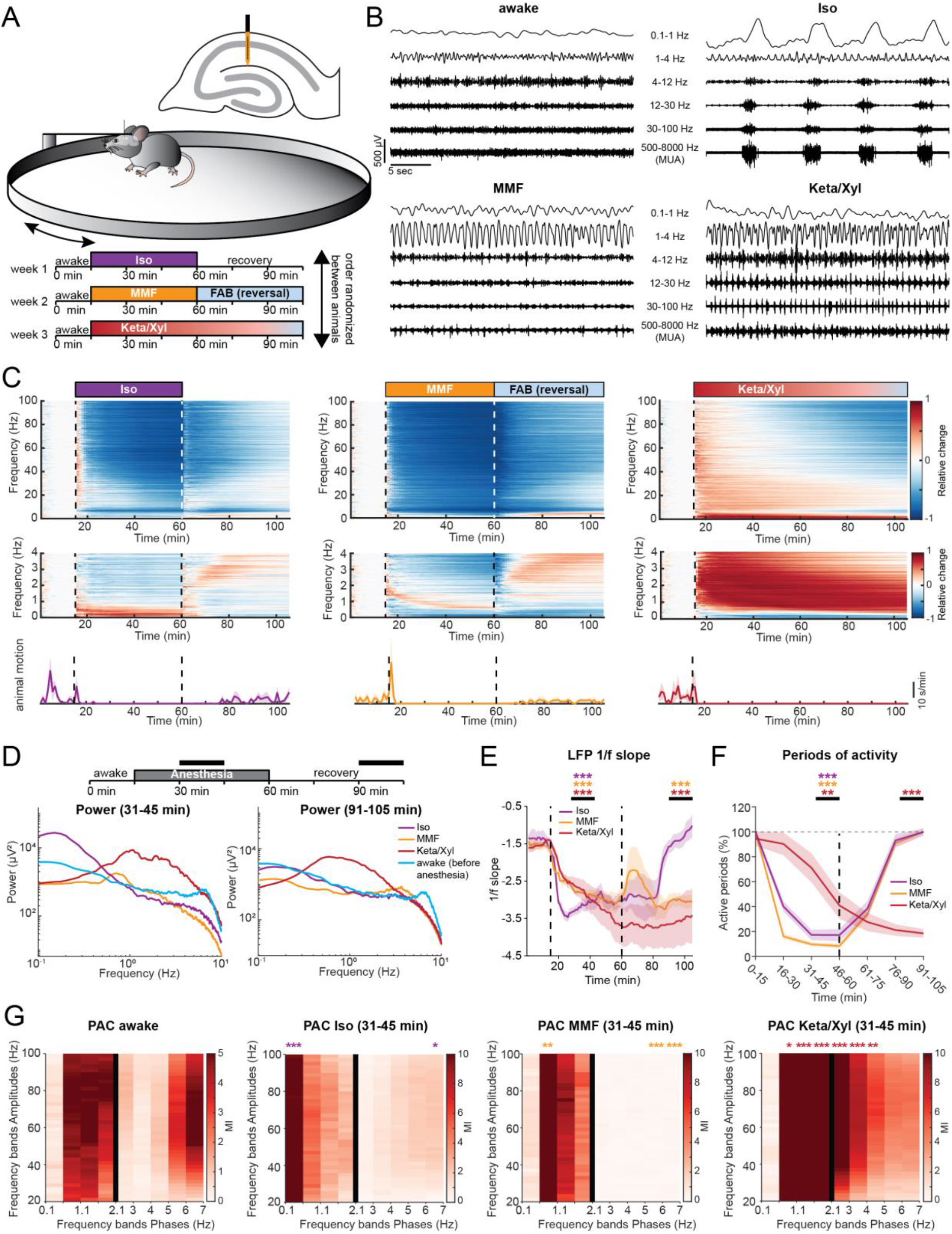
LFP recordings in dorsal CA1 during wakefulness and anesthesia reveal distinct and complex alterations by Iso, Keta/Xyl and MMF. **(A)** Experimental setup. Extracellular electrical recordings in dorsal CA1 were performed in four head-fixed mice for 105 min, continuously. Each animal was recorded under all anesthesia as indicated in the scheme. Order of anesthetics was pseud-randomized. **(B)** Characteristic local field potential (LFP) recordings during wakefulness and under three different anesthetics. **(C)** Color-coded modulation index (MI) plots (upper and middle panels) for LFP power and motion profiles (lower panels) for the three different anesthetic conditions. Upper panels display LFP power for 0-100 Hz frequency range, lower panels for 0-4 Hz. **(D)** Line plot displaying LFP power spectra for the two time periods indicated by horizontal black bars. For comparison, the 15-min spectrum of the awake period before anesthesia induction is plotted in both graphs. Statistical differences are indicated in Fig. S1C **(E)** Line plot displaying the power-law decay exponent (1/f) of the LFP power spectrum for the 30-50 Hz range. Lines display mean ± SEM. **(F)** Line plot displaying the fraction of active periods compared to the pre-anesthetic wakeful state, in 15 min bins throughout the entire recording duration. Lines display mean ± SEM. **(G)** Heat map displaying Phase-amplitude-coupling (PAC) for pre-anesthetic wakeful state (left) and for the indicated time periods during anesthesia. Different bin sizes (0.5 Hz and 1 Hz, separated by vertical black line) are used to resolve low- and high-frequency PAC. Vertical dashed lines in (C) and (E) indicate time points of anesthesia induction (Iso, MMF, Keta/Xyl) and reversal (Iso & MMF only). Vertical dashed line in (F) indicates time point of anesthesia reversal (Iso & MMF only). Asterisks in (E) and (F) indicate significance of time periods indicated by black horizontal line compared to 15-min period before anesthesia. Anesthetic conditions are color-coded. Asterisks in (G) indicate significant differences compared to the corresponding frequency band during wakefulness.^*^ p < 0.05, ^**^ p < 0.01, ^***^ p < 0.001, n = 4 mice. For full report of statistics, see statistics table.

It is widely accepted that, in the neocortex, GAs favor slow oscillations at the expense of faster ones [29]. To determine whether this is also the case in the hippocampus, we next asked how the investigated anesthetics affect slow network oscillations. Consistent with previous reports [30-32], Keta/Xyl strongly enhanced LFP power at 0.5-4 Hz throughout the entire recording period (Fig. 1C,D, S1C), but suppressed frequencies lower than 0.5 Hz. In contrast, Iso strongly augmented LFP power below 0.5 Hz, peaking at 0.1-0.2 Hz (Fig. 1C,D, S1C), whereas MMF induced no significant increase in the low-frequency regime. However, similar to Keta/Xyl, a significant reduction was present below 0.5 Hz, which persisted throughout the entire recording period (Fig. 1C,D). Analysis of the power-law decay exponent (1/f slope) of the LFP power spectrum facilitates detection of non-canonical changes in LFP power, including aperiodic (non-oscillatory) components [33]. The 1/f slope has been hypothesized to track excitation/inhibition (E/I) balance [34, 35], and is reduced in the cortex under anesthesia [36, 37], indicating a shift towards inhibition. Considering the robust effects on LFP power that we reported, we reasoned that the 1/f slope might also be altered. Indeed, all anesthetics significantly decreased the 1/f slope, albeit with a different temporal profile. While the effect of Iso occurred within a few minutes, MMF and Keta/Xyl operated on a longer timescale (Fig. 1E). Moreover, periods of activity were consistently and strongly reduced immediately under Iso and MMF, but delayed by 30 min under Keta/Xyl (Fig. 1F). These results indicate that all anesthetics shift the LFP to lower frequencies and tilt the E/I balance towards inhibition, albeit with different temporal profiles.

In contrast to Keta/Xyl-anesthesia, Iso- and MMF-anesthesia can be efficiently antagonized. Removing the face mask is sufficient to antagonize Iso-anesthesia, while antagonization of MMF-anesthesia requires injection of a wake-up cocktail (Flumazenil, Atipamezole and Buprenorphine, FAB) [38, 39]. 20-30 min after Iso withdrawal, animals regained motility and periods of silence in the LFP receded (Fig. 1C,F). However, in contrast to post-Iso, LFP power did not fully recover after FAB, remaining significantly reduced at frequencies below 0.5 and above 30 Hz for the entire 45 min-post anesthesia recording period (Fig. 1C,D). In contrast, elevated LFP power in the 0.5-4 Hz band and reduction in active periods remained significant throughout the entire recording in the presence of Keta/Xyl. In line with these results, the 1/f slope promptly reverted to values similar to baseline after Iso discontinuation. In contrast, the recovery was only transitory and partial after MMF antagonization, and virtually absent for Keta/Xyl (Fig. 1E), indicating that the E/I balance recovered only after Iso within 45 min.

Cross-frequency coupling between theta and gamma oscillations has been suggested to underlie information transfer in the hippocampus [40]. Given the strong decrease of theta power in the presence of Iso and MMF, we reasoned the phase modulation of the gamma rhythm could also be altered. To test this, we used phase-amplitude coupling (PAC) to measure whether the phase of slow LFP oscillations modulates the amplitude of the signal at a higher frequency. In line with previous results [41, 42] a significant coupling between theta and gamma frequency bands, as well as between frequencies in the 1-2 Hz range and gamma was present in the awake state (Fig. 1G). Moreover, anesthesia strongly altered PAC. In accordance with the LFP power analysis, the coupling reached a maximum strength between the dominant slow-frequency oscillations induced by the various anesthetics (<0.5 Hz for Iso, ∼1 Hz for MMF and 0.5-4 Hz for Keta/Xyl) and gamma (Fig. 1G). For all anesthetics, the range of phase-modulated amplitudes was wide, suggesting that the modulating phase corresponds to the identified slow-wave activity.

Taken together, these data show that all three GAs differently and persistently modulated the network oscillations in dCA1, a full recovery of activity being detected within 45 min only for Iso.

### Delayed recovery of neuronal spiking patterns after anesthesia

While the LFP provides information about general network states in the hippocampus, it is influenced by long-range activity and highly active regions in the vicinity of CA1 [43]. To assess the effects of GAs on CA1 neurons, we analyzed the spiking of individual units (56-72 units per animal, n=4 mice) before, during and after each of the anesthetic conditions. All anesthetics significantly and rapidly (<1 min) decreased spiking activity in CA1 neurons (Fig. 2A,B, S2), with MMF leading to the most potent suppression, followed by Iso and Keta/Xyl. These alterations were generally present in all layers of CA1 (Fig. S2C). Although the bulk spike rate was strongly reduced, the number of active neurons (see Methods) was only mildly affected (Fig. 2C), reaching a significant reduction only with MMF. This observation suggests that anesthesia broadly reduces neuronal activity, and does not modulate only a discrete subpopulation of neurons. Both firing rate and the number of active neurons recovered within 45 min after reversal for MMF and Iso (Fig. 2A-C, S2). As previously reported for NREM sleep [44], we found a negative correlation between the anesthesia-induced reduction of firing rate and the firing rate in wakefulness (Fig S2B).

**Figure 2:**
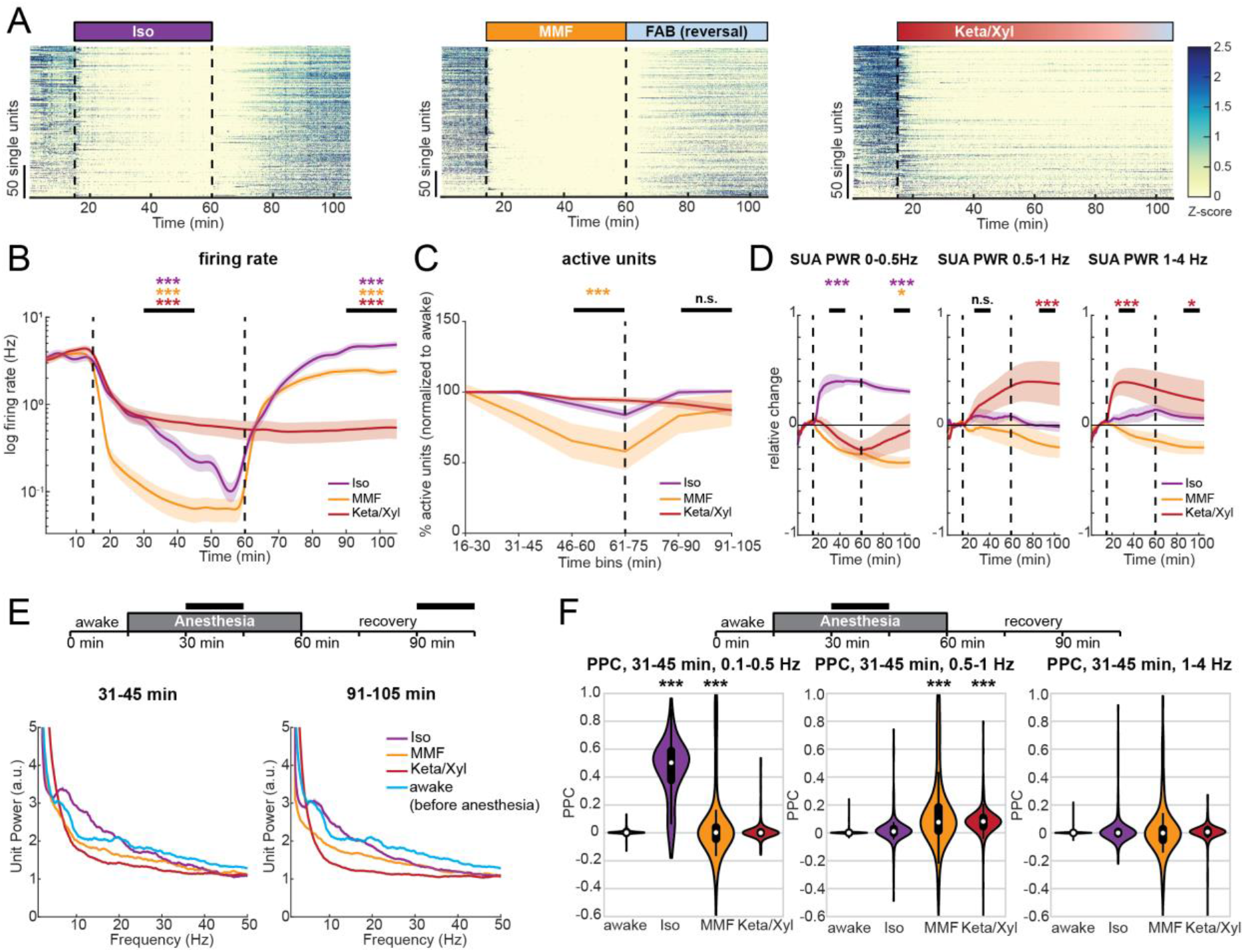
Single unit activity in dorsal CA1 is strongly reduced during anesthesia, and remains significantly altered long after its termination. **(A)** Raster plots of z-scored single-unit activity (SUA) for the three different anesthetic strategies in four mice. Units are sorted according to initial activity during wakefulness. **(B)** Line plot of SUA firing rate before, during and after anesthesia induction. **(C)** Line plot displaying the fraction of active units compared to the pre-anesthetic wakeful state, for all three anesthetics in 15 min bins throughout the entire recording duration. **(D)** Relative change of population firing rate power in the 0-0.5, 0.5-1 and 1-4 Hz frequency band. SUA PWR = power of SUA spike trains. **(E)** Line plot displaying the normalized power spectra of population firing rate for the two time periods indicated by horizontal black bars. For comparison, the 15-min spectrum for pre-anesthetic wakeful state is plotted in both graphs. **(F)** Pairwise phase consistency (PPC) at low frequencies in the same frequency bands as (D), for the indicated time points during anesthesia. White dots indicate median, vertical thick and thin lines indicate 1^st^-3^rd^ quartile and interquartile range, respectively. Colored lines in (B) - (D) display mean ± SEM. Vertical dashed lines in panels (A), (B) and (D) indicate time points of anesthesia induction (Iso, MMF, Keta/Xyl) and reversal (Iso & MMF only). The vertical dashed line in (C) indicates the time point of anesthesia reversal (Iso & MMF only). Asterisks in (B) - (D) indicate significance of periods indicated by black horizontal line compared to period before anesthesia. Anesthetic conditions are color-coded. Asterisks in (F) indicate significant differences to wakefulness. ^*^ p < 0.05, ^**^ p < 0.01,^***^ p < 0.001, n = 4 mice. For full report of statistics, see statistics table.

To investigate whether the rhythmicity of single neuron firing was affected similarly to the LFP, we analyzed the spectral properties of 1 ms-binned SUA firing (i.e., power of SUA spike trains, for details, see Methods). In the presence of Iso, SUA power was consistently increased in the range between 0 and 0.5 Hz (Fig. 2A,D, S2), in line with the strong modulation of LFP at 0.1-0.2 Hz. Of note, this effect did not vanish after Iso removal, suggesting that Iso has a long-lasting impact on firing rhythmicity. In contrast, and in line with its effects on the LFP, MMF generally reduced, albeit less strongly, SUA power, including the low frequencies. A significant reduction of SUA power was still present 45 min after antagonization in the 0-0.5 Hz band. Keta/Xyl, on the other hand, only showed a tendency towards reduced SUA power in the frequency band below 0.5 Hz, but increased SUA power significantly in the range between 0.5 and 4 Hz, consistent with its effect on the LFP (Fig. 2D). This modulation was present throughout the entire recording. At higher frequencies, Iso led to a peak in the theta frequency range, similar to wakefulness (Fig. 2E), yet it reduced the SUA power in the beta/gamma range. Keta/Xyl and MMF caused an overall reduction in SUA power at frequencies >5 Hz (Fig. 2E). Thus, GAs differentially impair spiking rhythmicity. These changes appeared to follow similar dynamics than those in the LFP.

To confirm the synchrony between spikes and low-frequency oscillations, we calculated their pairwise phase consistency (PPC) [45]. When compared to pre-anesthesia, PPC values for the 0.1-0.5 Hz frequency band were augmented by Iso. Keta/Xyl increased coupling of spikes to the LFP between 0.5 and 1 Hz, whereas MMF showed a weak, but significant increase of coupling at frequencies below 1 Hz (Fig. 2F).

Similar to the LFP, the SUA firing rate nearly fully recovered during the 45 min post-Iso (Fig. 2A,B, S2), with even a slight, but significant increase at the end of the recording period. In contrast, after FAB-induced MMF reversal, CA1 spiking activity remained slightly reduced, reflecting the lack of LFP recovery. For Keta/Xyl, SUA remained suppressed during the entire recording period (Fig. 2B). Strikingly, SUA power did not fully recover for any of the tested anesthetics (Fig. 2E).

Taken together, we show that all investigated GAs caused a persistent and robust reduction of CA1 firing. Moreover, spiking during anesthesia was phase-locked to the GA-induced slow network oscillations.

### Iso, Keta/Xyl and MMF reduce number, amplitude, and duration of calcium transients

To monitor the population dynamics of CA1 neurons in the presence of different anesthetics, we imaged the same field of view (FOV) using the genetically encoded indicator GCaMP6f [46] and systematically compared the activity of identified neurons during quiet wakefulness and in the presence of different anesthetics (Fig. 3A).

**Figure 3:**
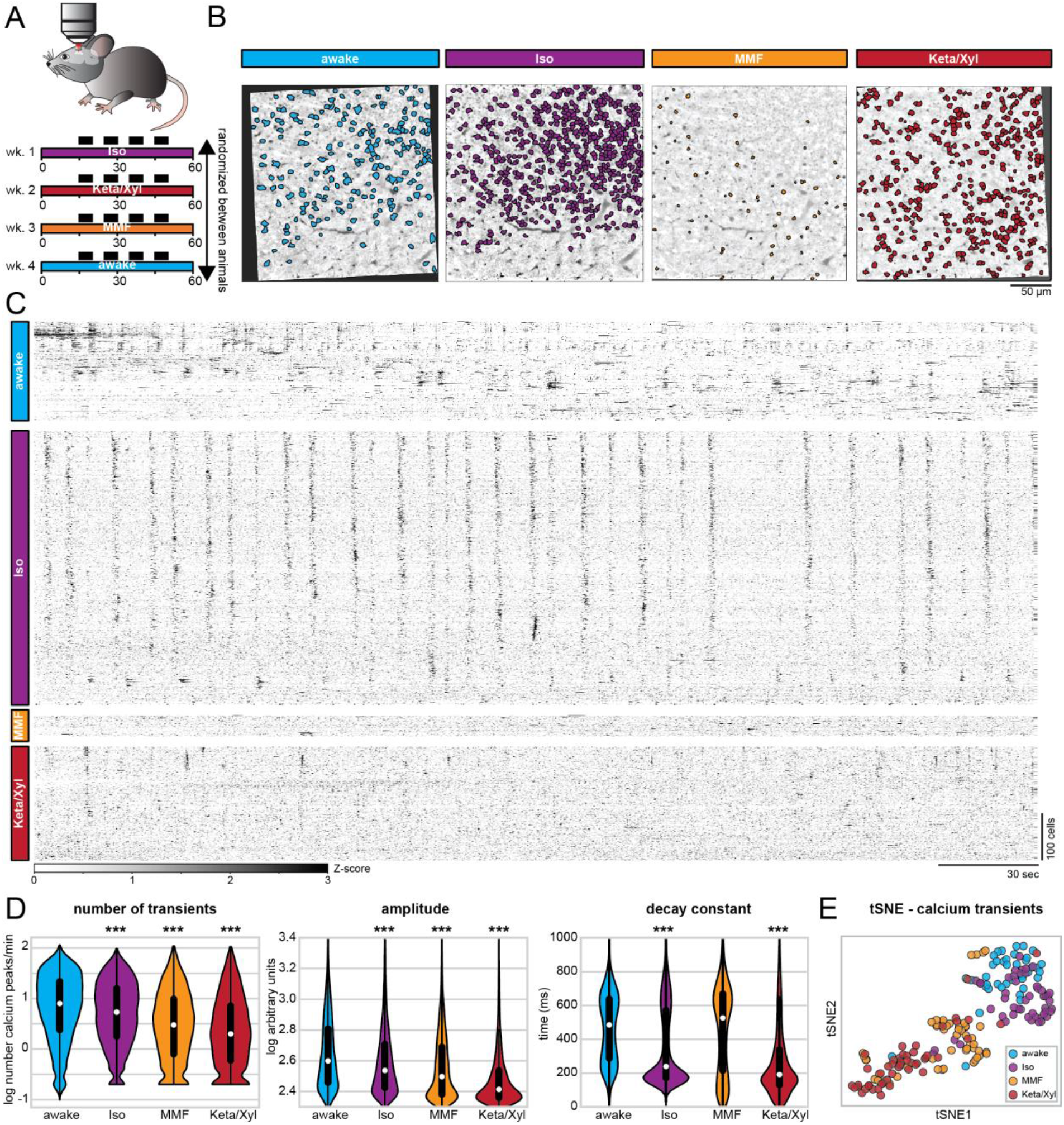
Repeated calcium imaging in dorsal CA1 reveals distinct activity profiles for Iso, MMF and Keta/Xyl. **(A)** Experimental strategy for chronic calcium imaging of cellular activity in dorsal CA1. For each condition, seven mice were imaged four times for five minutes, as indicated by black fields in the scheme. The order of imaging conditions was pseudo-randomized. **(B)** Time-averaged, two-photon images of the same FOV in CA1 aligned to the Iso condition. ROIs of automatically extracted, active neurons are overlaid for each condition. **(C)** Raster plots of z-scored calcium transients in the same animal under different conditions. Traces are sorted by similarity. **(D)** Violin plots quantifying the number (left), amplitude (middle), and decay (right) of detected calcium transients. White dots indicate median, vertical thick and thin lines indicate 1^st^-3^rd^ quartile and interquartile range, respectively. **(E)** tSNE plot summarizing the average calcium transients properties. Each data point represents one recording session. Asterisks in (D) indicate significant differences to wakefulness. *** p < 0.001. Note, to facilitate readability, only differences to wakefulness are indicated. For full report of statistics, see statistics table.

First, we considered all active neurons in each condition and analyzed the average rate (i.e., the number of transients), amplitude, and duration (i.e., the decay constant) of calcium transients across all imaging sessions in 7 mice. In line with the results of SUA analysis (see Fig. 2C), a large number of CA1 pyramidal neurons were active in the presence of all three GAs. Using extraction parameters that restricted the number of ROIs but maximized signal quality (see Methods), we obtained a median of 311 (min-max of 16-817) active neurons per FOV, for a total of 189 five-minutes recordings. All GAs significantly altered calcium dynamics in CA1 neurons, reducing the activity (Fig. 3C,D), as previously shown for neuronal spiking (Fig. 2B). Also, in line with the effect on SUA (Fig. S2B), the magnitude of the anesthesia-induced reduction of calcium transients was negatively correlated with the wakefulness calcium transients rate (Fig. S6D). However, each condition could be characterized by a specific signature in their calcium dynamics. Iso yielded only a mild decrease of rate and amplitude, but a strong reduction of duration of calcium transients (Fig. 3D). Consistent with effects on LFP and SUA, calcium transients showed a spectral peak between 0.1 and 0.2 Hz (Fig. S4). In contrast to Iso, MMF did not significantly affect the duration of transients but reduced their rate and amplitude when compared to wakefulness. Keta/Xyl-anesthesia had the strongest effect on calcium transients, leading to a reduction of all three parameters compared to wakefulness (Fig. 3D). Unlike for electrophysiological recordings, no spectral peak was present in calcium transients, most likely due to the strong suppression of calcium activity by Keta/Xyl. Considering all parameters, the four groups tended to segregate into clusters, one consisting mostly of recordings under Keta/Xyl, and another one consisting of awake and Iso recordings. Most recordings under MMF clustered between these two groups (Fig. 3E). Importantly, these findings were robust to changes in the signal extraction pipeline. Varying the threshold for calcium transient detection across a wide range of values did not affect the reported effects on rate and height of transients (Fig. S3B). Further, conducting the same analysis on neuronal activity metrics that are independent of calcium transients detection (integral and standard deviation) or on dF/F calcium signals also yielded analogous results (Fig. S3C-E).

### Iso, Keta/Xyl and MMF distinctly modulate cellular calcium dynamics in individual neurons

One possible explanation for these distinct modes of calcium activity could be that each anesthetic condition recruits a unique set of neurons characterized by particular spiking properties. We tested this possibility by analyzing calcium transients in neurons that were active during all conditions (Fig. 4A, S5, S6). To obtain a sufficient number of active neurons, we extracted calcium transients using a lower quality threshold, accepting more neurons per recording (see Methods). In this manner, we obtained a median of 783 neurons per recording (min-max of 156-1641). While this shifted the overall distribution of calcium parameters to lower values, the relative ratios between the four conditions remained the same and the differences between anesthesia groups were preserved (Fig. S3F-G). Also, when considering only neurons that were active in all four conditions, rate as well as amplitude of calcium peaks were generally reduced under anesthesia, being lowest in the Keta/Xyl condition (Fig. 4B,C). Compared to the whole dataset, differences in decay constant were less pronounced. The median decay constant strongly decreased for awake and MMF conditions, while it increased for Iso and Keta/Xyl. These results indicate that both the between-as well as the within-condition variance strongly decreased when considering only neurons active under all conditions.

**Figure 4:**
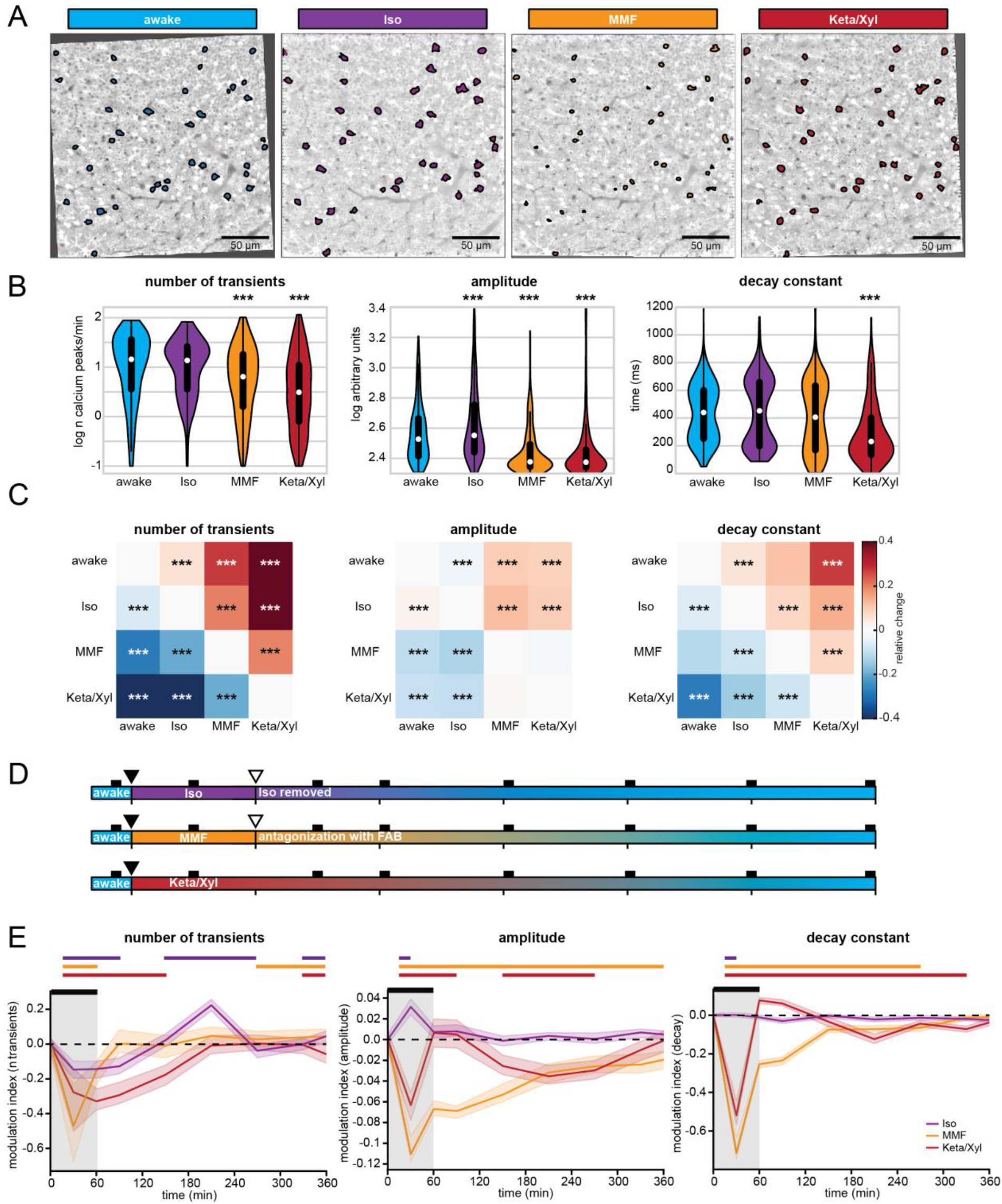
Calcium activity profiles in neurons active during all conditions are similar between wakefulness and Iso. **(A)** Two-photon time-averaged images of the same FOV in CA1, aligned to the Iso condition (same images as in figs. 3). ROIs show neurons active in each condition, allowing direct comparison of calcium transients in the same cells under different conditions. **(B)** Violin plots quantifying the number (left), amplitude (middle), and decay (right) of detected calcium transients. White dots indicate median, vertical thick and thin lines indicate 1^st^-3^rd^ quartile and interquartile range, respectively.**(C)** Heat maps displaying the relative change in the number (left), amplitude (middle), and decay (right) of calcium transients between neurons active in pairs of conditions (see also Fig. S6C). **(D)** Schematic representation of long-term calcium imaging experiments to assess recovery from anesthesia. Black rectangles indicate imaging time points (up to 10 min duration each). Filled and open triangles indicate the start and end of the anesthesia period. **(E)** Line diagrams showing the relative change (modulationindex) of the median number of calcium transients (left), their amplitude (middle), and decay constant (right) during anesthesia and recovery relative to the awake state before anesthesia induction. The black bar indicates the anesthesia period. Shaded, colored lines indicate 95% confidence interval. Note, Keta/Xyl anesthesia could not be terminated. The horizontal, colored lines indicate significant difference (p < 0.05) to awake time point (t = 0) for the respective condition. Asterisks in (B) and (C) indicate significant differences to wakefulness. ^***^ p < 0.001. Note, to facilitate readability, only differences to wakefulness are indicated. For full report of statistics, see statistics table.

The relatively low number of neurons active in all four conditions (335 neurons) limited the statistical analysis. Therefore, we compared neurons that were active in any two combinations of conditions (Fig. S6C). This analysis further corroborated the similarity of neurons active during wakefulness and Iso anesthesia (Fig. 4C, S6C). Rate, amplitude, and duration of calcium transients were most similar between wakefulness and Iso compared to the other GAs. In contrast, neurons active during wakefulness and either Keta/Xyl or MMF showed decreased rate, amplitude and duration under anesthesia, with Keta/Xyl causing the strongest phenotype (Fig S6C). Overall, this indicates that anesthetics influence the firing properties of hippocampal neurons. However, the magnitude and direction of these effects vary considerably. Iso anesthesia has the mildest effect, and it most likely arises from distinct neuronal populations being active in the two conditions (wakefulness vs. Iso anesthesia), as the firing properties of cells that are active in both are barely affected (Fig. 4B,C). On the other hand, the strong effects of MMF and Keta/Xyl on all calcium parameters in the same cells indicate that different anesthetics directly alter the firing properties of individual neurons. Thus, alterations in firing properties of neuronal populations (e.g., SUA, Fig. 2B-D) are not solely explainable by different subpopulations of neurons being active between awake and anesthesia.

### Population activity recovers with different temporal dynamics after Iso, Keta/Xyl and MMF

The LFP recordings showed that network activity remained altered for 1.5 h after Keta/Xyl injection, but also after antagonization of MMF, while most aspects returned to pre-anesthetic conditions during 45 min after Iso removal. To assess network effects of the different anesthetics on a longer time scale, we used repeated calcium imaging during 6 hours after anesthesia onset and 5 hours after Iso termination and MMF antagonization (Fig. 4D). In line with our previous results, the number of calcium transients was strongly reduced 30 min after MMF or Keta/Xyl injection, while the reduction had a lower magnitude for Iso. Similarly, MMF and Keta/Xyl most strongly reduced the amplitude and duration of calcium transients, while Iso mildly increased amplitude without affecting the decay constant (Fig. 4E).

Confirming the action dynamics monitored by LFP recordings in vivo, recovery from Iso anesthesia was fast and only the rate of transients mildly changed during the hours after removing the mask. In contrast, after Keta/Xyl injection, amplitude and duration of transients were altered throughout the following 6 hours, while the reduction of the calcium transients rate was not reverted until up to 4 hours later. Recovery to the pre-anesthetic state was even slower after MMF/FAB. Despite antagonization of MMF anesthesia with FAB, calcium transients remained disturbed for up to 6 hours. Thus, the different anesthetics not only induce unique alterations of CA1 network dynamics, but also show different recovery profiles (Fig. S6E).

### Anesthesia decorrelates hippocampal activity

Calcium imaging studies in the visual cortex of ketamine anesthetized rats [27] and Iso anesthetized mice [25] showed that anesthesia increases the overall pairwise correlations between firing neurons and, consequently, induces more structured patterns of activity. While neocortical L2/3 cells typically show a high degree of local interconnectivity [47], this is not the case for CA1, where pyramidal cells receive their main excitatory input from CA3 and entorhinal cortex and send their efferents to subiculum and extrahippocampal areas [9]. Another difference between neocortex and hippocampal CA1 area is that the neocortex receives strong direct input from primary thalamus, which is a major source for slow oscillations during anesthesia-induced unconsciousness and sleep [1, 48, 49]. In comparison to neocortex, hippocampus shows different patterns of activity, including sharp waves, which are generated intrinsically in the hippocampus, likely originating in CA3 [50]. To investigate whether these differences cause a different impact of anesthesia on the population activity in CA1 when compared to the neocortex, we analyzed the dynamical structure of population activity using both calcium imaging and SUA of extracellular recordings in vivo. First, we analyzed Fisher-corrected Pearson pairwise correlation between neuropil-corrected raw fluorescence traces. We found that both correlation and anticorrelation were highest in animals during quiet wakefulness (Fig 5A-B). In particular, the awake condition had a higher proportion of correlation coefficients both in the 1^st^ as well as in the 4^th^ quartile of the entire distribution and, accordingly, higher absolute correlation values (Fig. 5B, S7A). Similar to the firing properties (SUA, fig. 2), Iso induced the milder changes, whereas Keta/Xyl caused the strongest phenotype. This relationship was preserved in neurons active during all conditions (Fig. S7B), indicating that anesthesia generally reduces correlated activity between neurons and that this effect is not attributable to the activity of particular neuronal subpopulations. Moreover, these effects were not influenced by the distance between the pair of neurons whose correlation was quantified (Fig. 5C). These findings highlight the major differences between the anesthesia-induced effects on neuronal coupling in hippocampal CA1 and neocortex. In accordance with the anatomy of CA1, the correlation between pairs of neurons was only mildly affected by the distance between them, with or without anesthesia. Not only were neurons less highly correlated to each other under anesthesia, but their coupling to the whole population activity [51] was reduced as well. The proportion of neurons with population coupling in the 4^th^ quartile of the entire distribution was highest for awake, and most strongly reduced under Keta/Xyl and MMF, while Iso showed only mild effects (Fig. 5D).

**Fig 5.**
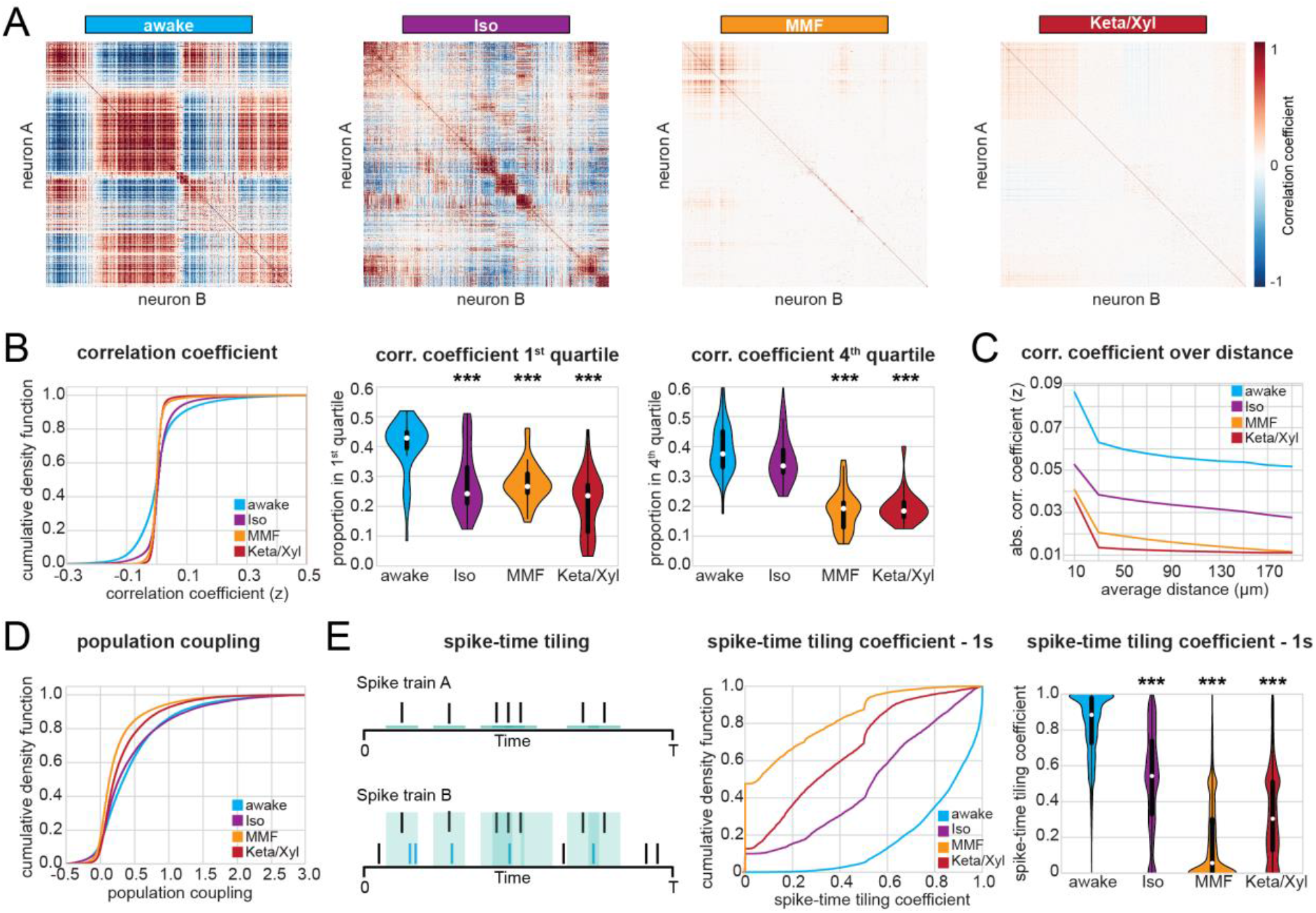
Correlation analysis of CA1 calcium activity and SUA shows decorrelation under anesthesia. **(A)** Heat maps displaying representative correlation matrices of calcium activity between pairs of neurons during wakefulness and the three different anesthetic conditions in the same animal. Matrices are sorted by similarity. **(B)** Left: Line plot displaying cumulative distribution of Fisher-corrected Pearson correlation coefficients between pairs of neurons (calcium imaging). Center: violin plot displaying the proportion of pairs found in the 1^st^ (most negative) and 4^th^ (most positive) quartile of the distribution. **(C)** Line plot displaying the absolute pairwise correlation coefficients over distance (calcium imaging, 25 micrometer bins). **(D)** Line plot displaying the cumulative distribution of population coupling (calcium imaging). **(E)** Quantification of correlation between pairs of extracellularly recorded single units using the spike-time tiling coefficient (STTC). Left: Schematic illustration of the STTC quantification. Center: cumulative distribution of the STTC with a 1000 ms integration window. Right: violin plot quantifying the STTC. In violin plots, white dots indicate median, vertical thick and thin lines indicate 1^st^-3^rd^ quartile and interquartile range, respectively. Asterisks in (B) and (E) indicate significant differences to wakefulness. ^***^ p < 0.001. Note, only differences to wakefulness are indicated. For comparison between conditions, see statistics table.

To further relate the calcium imaging data to extracellular recordings of neuronal firing, we carried out an analogous analysis on SUA. To avoid the confounding effect of firing rate, we quantified the correlation between pairs of neurons using the spike-time tiling coefficient [52], a measure that is largely insensitive to variations of the firing rate (see Methods). To be consistent with the calcium data, we quantified correlations within 1 second, a timescale of the same magnitude as the decay constant used to extract calcium signals (700 ms). This analysis confirmed that all anesthetics decorrelated neuronal activity (Fig. 5E). This effect was still present, albeit less pronounced, using an integration window of 10 ms, which is closer to the duration of action potentials (Fig. S7C). Overall, the decorrelation was milder under Iso anesthesia and stronger under Keta/Xyl and MMF. Thus, all three GAs decorrelated calcium transients and spiking activity in the CA1 area, with MMF and Keta/Xyl inducing the most prominent effects.

### Anesthesia fragments temporal and spatial structure of hippocampal activity

The decorrelation of neuronal activity during anesthesia suggests that GAs might impact the spatial and temporal organization of CA1 neuronal ensembles (see Fig. 5A). To test this hypothesis, we analyzed the same number of active neurons for each condition, since a different number of neurons in each condition potentially influences the number and size of detected clusters [26]. First, we monitored the impact of GAs on the temporal structure of CA1 activity. We defined the number of clusters identified by principal component analysis (PCA) as the number of components that were needed to explain 90% of the variance. Moreover, we assessed the power-law slope of variance explained over the first 150 components (Fig. 6A). Both methods led to a larger number of clusters and a flatter power-law slope for anesthesia when compared to wakefulness (Fig. 6A). Further corroborating these findings, both tSNE dimensionality reduction and affinity propagation (AP) clustering (see Methods) also revealed a larger number of clusters for anesthesia compared to wakefulness (Fig. 6B,C). These observations indicate that activity is less structured under anesthesia. In line with previous results, Iso had the weakest effect, whereas Keta/Xyl consistently induced the most pronounced phenotype. Analysis of the deconvolved calcium traces led to comparable results (Fig S8A,B). These findings support the idea that GAs fragment the hippocampal network into a more diverse repertoire of microstates.

**Fig 6.**
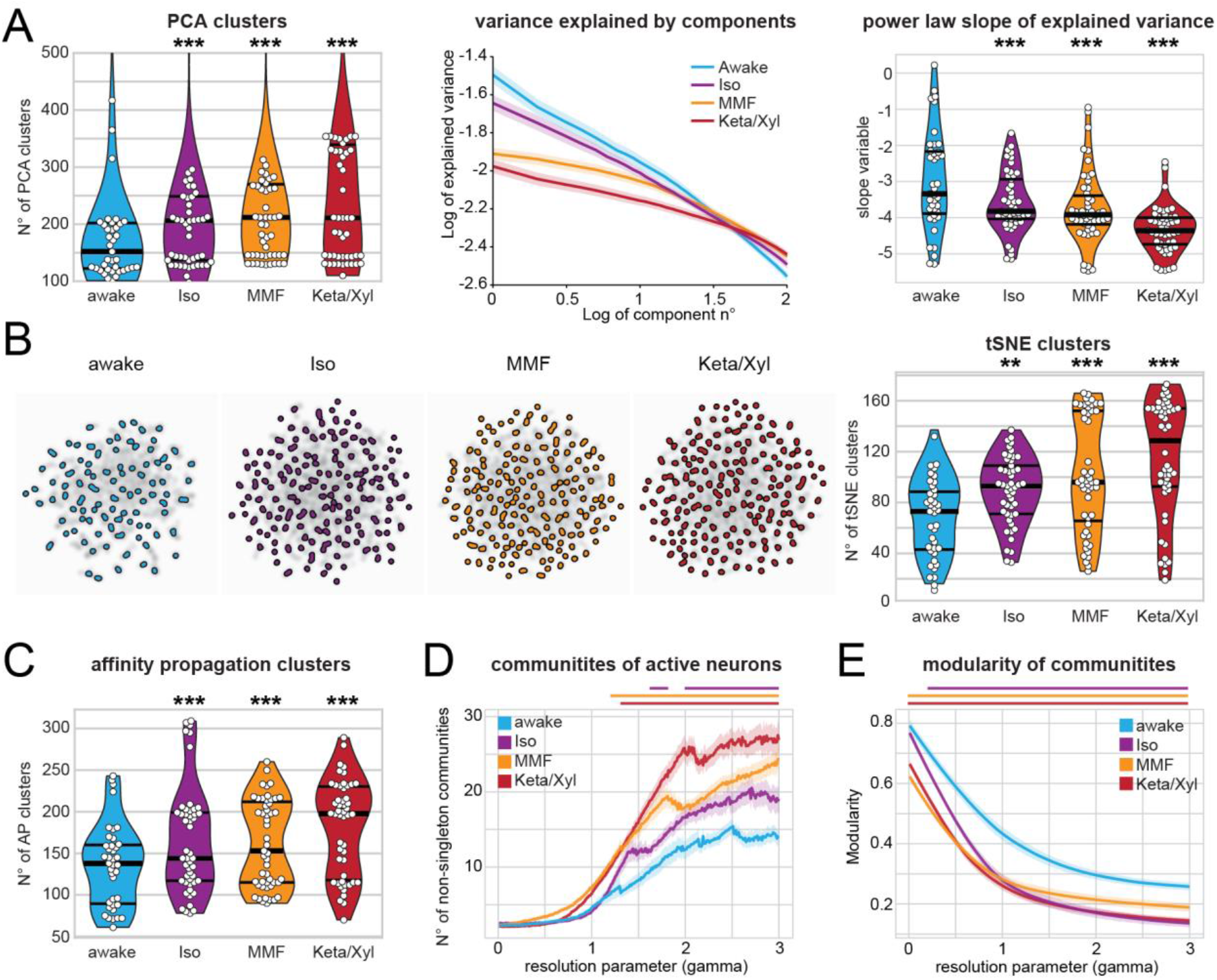
Calcium activity in CA1 is temporally and spatially fragmented during anesthesia. **(A)** Left: violin plot quantifying the number of principal component analysis (PCA) clusters during wakefulness or anesthesia, as indicated. Middle: log-log line plot displaying the variance explained by the first 100 components for each condition. Right: violin plot quantifying the power-law slope of the variance explained by the first 100 components for each condition. **(B)** Left: tSNE plots of network events recorded in the same animal under the four indicated conditions. Right: Violin plot quantifying the number of tSNE clusters obtained from calcium recordings during the four different treatments. **(C)** Violin plot quantifying the number of clusters obtained by affinity propagation from calcium recordings during the four different treatments. **(D)** and **(E)** Line plots quantifying the number of detected communities and the modularity of the detected communities with the resolution parameter gamma ranging from 0 to 3. Horizontal lines in violin plots indicate median and 1^st^-3^rd^ quartile. Asterisks in (A) - (C) indicate significant differences to wakefulness. ^**^ p < 0.01, ^***^ p < 0.001. Horizontal lines above plots in (D) - (E) indicate significant difference to wakefulness. Anesthetic conditions are color-coded. Note, only differences to wakefulness are indicated. For comparison between conditions, see statistics table.

Second, we tested whether anesthesia disrupted the spatial structure of hippocampal activity, employing a modularity maximization approach [53, 54] designed to detect internally densely connected communities (modules). To allow detection of modules at varying sizes, we carried out our analysis while varying a resolution parameter (gamma) and thus focusing on different spatial scales. Using this approach, we showed that GAs increase the number of detected communities over a wide range of resolution parameter values (Fig 6D). Moreover, the modularity of these communities was lower than in wakefulness (Fig 6E). These results indicate that anesthesia results in a more fractured network with, on average, smaller and less coherent communities. A multi-resolution approach [55] followed by the selection of partitions based on hierarchical consensus clustering yielded similar results (Fig. S8C). Among GAs, Iso induced the mildest phenotype, whereas Keta/Xyl had the most prominent effects. Thus, GAs not only decorrelate hippocampal activity, but also consistently fragment both its temporal and spatial structure.

### Network alterations during sleep are less pronounced compared to anesthesia

Altered CA1 activity under anesthesia may affect synaptic function and memory processing. A naturally-occurring form of unconsciousness is sleep, which is required for network processes involved in memory consolidation [49, 56]. To decide whether the network perturbations described above resemble those naturally occurring during sleep, we first monitored CA1 activity by recording the LFP and spiking together with animal motion and the neck-muscle electro-myogram (EMG) in head fixed mice (Fig. S9A). We classified the signal into 30 s-long epochs of wake, rapid-eye-movement (REM) and non-REM (NREM) sleep. Further, a certain fraction of epochs, which we labelled as “uncertain”, could not be reliably classified into any of the previous three categories, (see Methods for details). Given that the behavioral attribution of these epochs is uncertain and difficult to interpret, we excluded them from further analysis. The animals spent most of their sleeping time in the NREM phase, with only short periods of intermittent REM sleep (Fig. 7A,I). The LFP showed enhanced theta power during REM phases, while the power at low frequencies was broadly increased during NREM sleep (Fig. 7B,C). Compared to anesthesia (Fig. 1), these changes in the LFP both during REM and NREM phases were modest. Along the same line, 1/f slope during NREM and REM sleep slightly decreased, indicating a small reduction of the E/I balance that had a significantly lower magnitude than the perturbation induced by GAs (Fig. 1D). Furthermore, the SUA rate was slightly reduced (Fig. 7E-F), in contrast to all anesthetics, that strongly suppressed firing (Fig. 2). As previously reported [44], and similarly to the effect of GAs, we detected a small but significant negative correlation between the NREM-induced reduction of firing rate and the wakefulness firing rate, whereas the effect failed to reach statistical significance for REM sleep alone (Fig. 7G). Moreover, NREM sleep induced a small reduction of pairwise correlation between pairs of neurons, as measured by the spike-time tiling coefficient with an integration window of one second.

**Fig 7.**
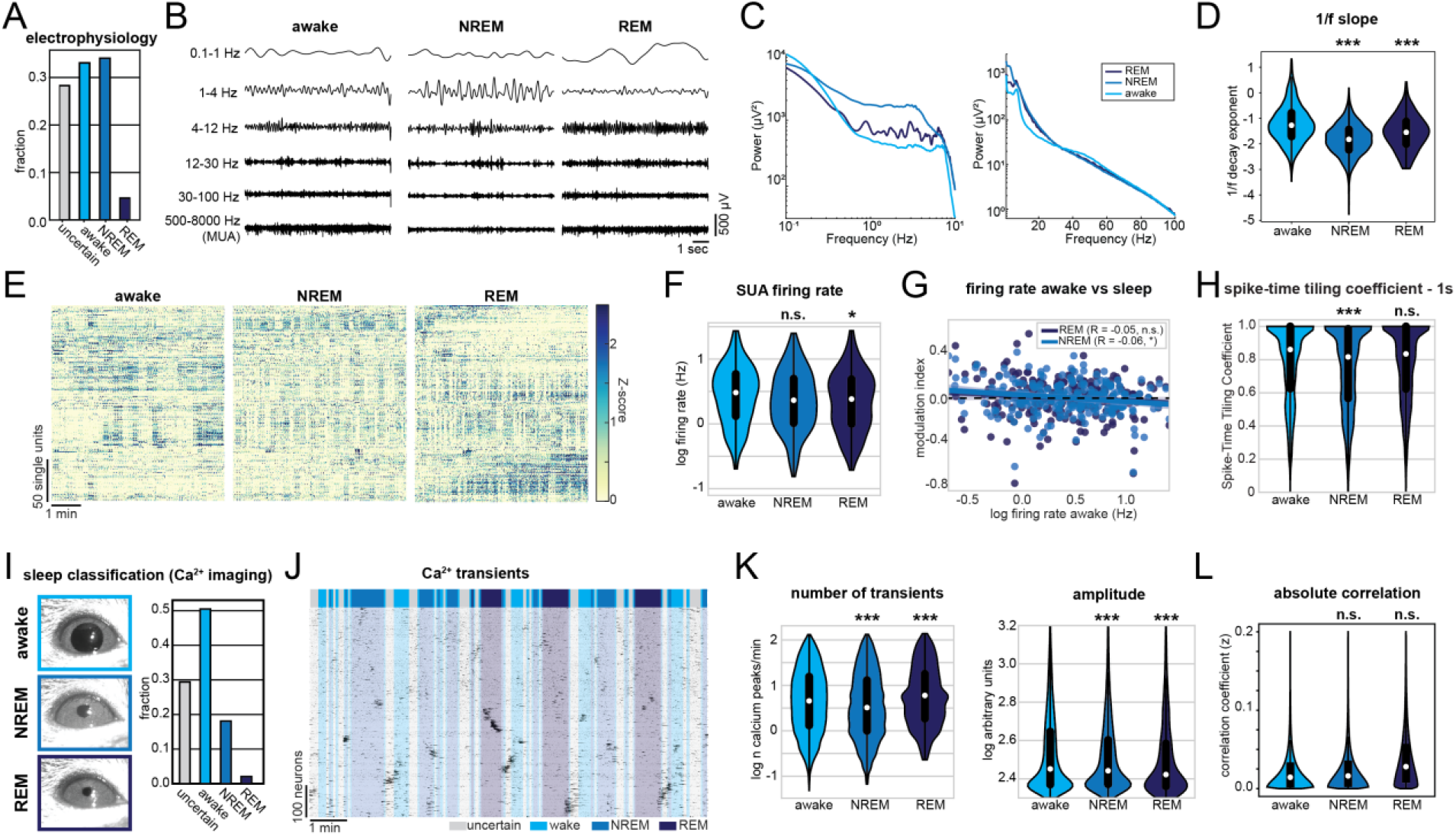
Sleep alters CA1 activity in a similar way to anesthesia but with a lower magnitude. **(A)** Classification of activity states during electrical recordings. **(B)** Characteristic LFP recordings during wakefulness, NREM and REM sleep. **(C)** Line plot displaying LFP power spectra for the indicated activity states. **(D)** Violin plot displaying the power-law decay exponent (1/f) of the LFP power spectrum. **(E)** Raster plots of z-scored single-unit activity (SUA) for the three different activity states in four mice. Units are sorted according to initial activity during wakefulness. **(F)** Violin plot showing SUA firing rate. **(G)** Scatter plot showing modulation of SUA firing rate during NREM (light blue) and REM sleep (dark blue) with respect to activity during wakefulness. **(H)** Violin plot quantifying the STTC. **(I)** Classification of activity states during CA1 calcium imaging based on eye videography. **(J)** Raster plots of z-scored calcium transients in an example recording of one animal transiting between wakefulness and sleep. Traces are sorted by similarity. **(K)** Violin plots quantifying the number (left), and amplitude (right) of detected calcium transients. **(L)** Violin plots quantifying absolute pairwise correlation of all recorded neurons. White dots indicate median, vertical thick and thin lines indicate 1^st^-3^rd^ quartile and interquartile range, respectively. ^*^ p < 0.05, ^**^ p < 0.01, ^***^ p < 0.001 w.r.t. to wake state, n = 3-7 mice.

To additionally investigate the effect of sleep on hippocampal activity, we used the above-mentioned recordings to train a machine-learning algorithm to classify wakefulness, NREM and REM sleep from eye videography images alone (Fig. S9) [57] (see Methods for details). In line with previous results, we were able to reliably distinguish wakefulness and NREM sleep (4-fold cross-validation accuracy >85%), whereas REM classification was less precise (4-fold cross-validation accuracy ∼30%). This classifier was then used to predict the physiological state of mice from which we recorded calcium transients in CA1 neurons. In the calcium imaging dataset, sleep was dominated by the NREM phase and only 17 min of REM sleep could be detected in a total of 864 min (Fig. 7I). Given the limited amount of detected REM sleep, its effects on hippocampal calcium activity should be interpreted with caution. As reported for LFP data, NREM only mildly reduced the rate of calcium transients, whereas REM sleep induced a small increase. In contrast, both NREM and REM sleep caused a small reduction in transient amplitude (Fig. 7K). Further, we did not detect an effect of the sleep state on absolute pairwise correlations (Fig. 7L).

In conclusion, sleep and GAs similarly affect the CA1 activity. However, the magnitude of effects was much smaller for sleep than for GAs. Both NREM and REM states were more similar to wakefulness than to the anesthetic state. Compared to the three different anesthetics, sleep had the closest resemblance to Iso. Thus, among the three different anesthetics, network alterations under Iso deviate the least from natural states such as wakefulness and sleep.

### Repeated anesthesia alters spine dynamics in CA1

The impact of Iso, MMF, and Keta/Xyl on CA1 activity might alter spine dynamics at CA1 pyramidal neurons. This issue is of critical relevance, since GAs disrupt activity patterns during development [58] also involving alteration of synaptic connectivity [59-61], but less is known about the impact of GAs on hippocampal synaptic structure during adulthood. So far, spine dynamics in hippocampus were only investigated under anesthesia, lacking comparison to the wake state. Moreover, the reported turnover rates varied strongly between studies [21, 62, 63]. Thus, it is unknown how repeated anesthesia in itself affects spine stability.

We repeatedly imaged the same basal, oblique, and tuft dendritic segments of CA1 pyramidal neurons under all four conditions (five times per condition, every four days), interrupted by a 30-day recovery period between conditions (Fig. 8A, S10A). To rule out time-effects, we pseudo-randomized the order of anesthetics (Fig. S10A). During wakefulness, without any anesthesia in between, the turnover ratio of spines on all dendrites was on average 18.6 - 20.5 % per four days. This turnover ratio was stable and did not change systematically over successive imaging sessions (Fig. 8B). Notably, all anesthetics affected spine turnover. Both MMF and Iso anesthesia mildly increased the turnover ratio compared to wakefulness (21.1 – 23.8 % for MMF, 24.0 – 24.7 % for Iso). Iso did not alter the surviving fraction of spines. Together with the significant increase in spine density over time (Fig. 8B) these results indicate that the elevated turnover ratio was due to a rise in the gained fraction of spines (Fig. S10B). In contrast, MMF led to a slight increase in the fraction of lost spines (Fig. S10B) and correspondingly, slightly decreased the surviving fraction compared to wakefulness. Spine density did not change over time. Keta/Xyl anesthesia showed the strongest effect on spine turnover (13.4 - 15.7 %), which was opposite to MMF and Iso, and therefore significantly lower rather than higher compared to the awake condition (Fig. 8B). This lower turnover ratio was accompanied by a higher surviving fraction and an increase in density with time (Fig. 8B). Consistently, the fraction of lost spines was most strongly reduced (Fig. S10B). Thus, Keta/Xyl anesthesia resulted in marked stabilization of existing spines and a reduction in the formation of new spines, indicative of a significant effect on structural plasticity. These effects were present on basal dendrites of stratum oriens (S.O.), oblique dendrites in stratum radiatum (S.R.) and tuft dendrites in stratum lacunosum moleculare (S.L.M.), albeit with different magnitude. Under Keta/Xyl the strongest impact on spine density was present in S.L.M., while turnover was most strongly reduced in S.O. Also, the increased spine turnover seen under Iso and MMF was most pronounced in S.O. (Fig. S10D).

**Fig 8.**
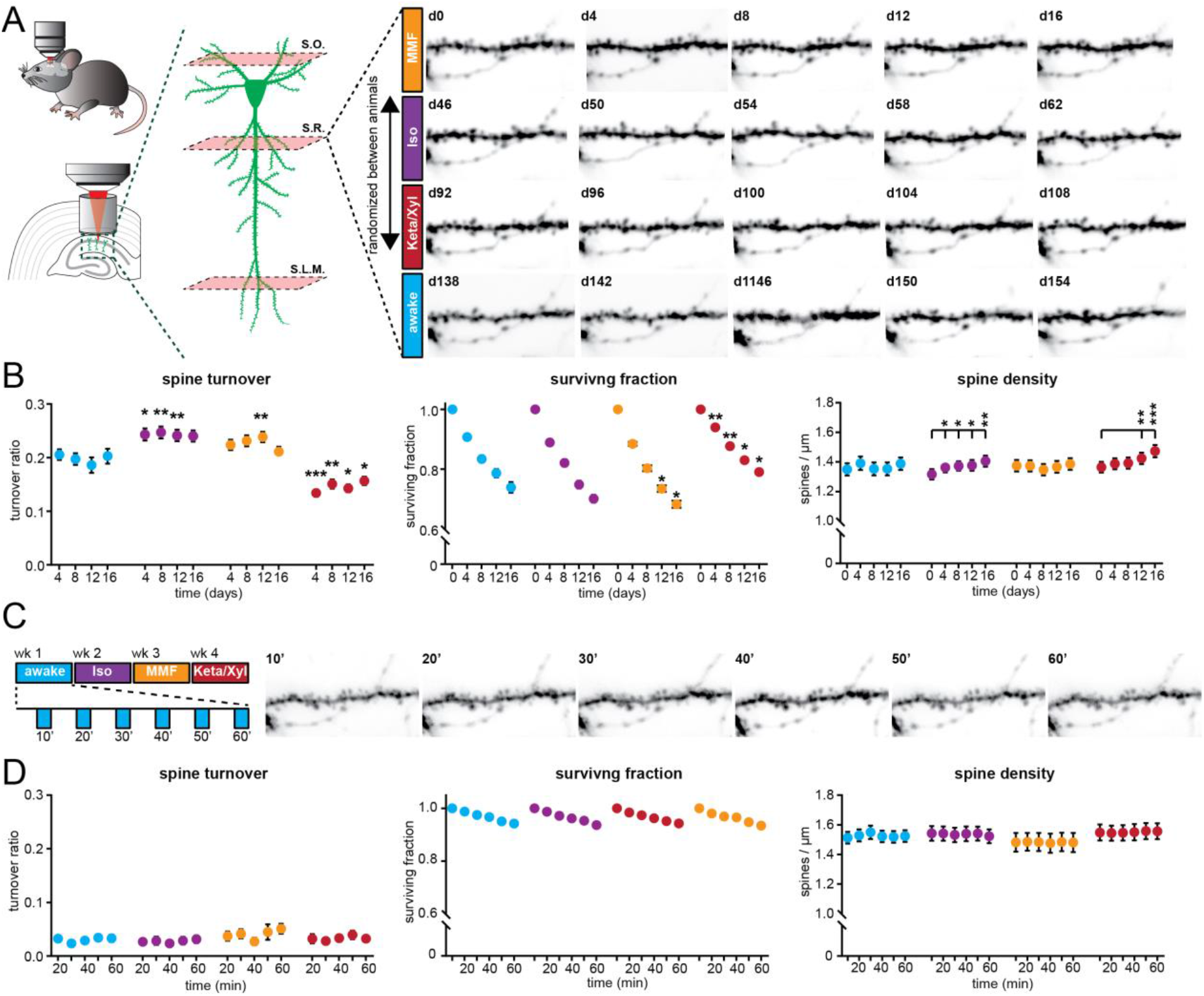
Spine turnover at CA1 pyramidal neurons is distinctly altered by repeated application of Iso, MMF and Keta/Xyl. **(A)** Left: Schematic illustration of in vivo spine imaging strategy. In each animal, spines were imaged on basal dendrites located in stratum oriens (S.O.), oblique dendrites in stratum radiatum (S.R.) and tuft dendrites in stratum lacunosum moleculare (S.L.M.). Right: Example showing an oblique dendrite in S.R. imaged chronically during all conditions. The order of anesthetic treatments was pseudo-randomized between mice (see Fig. S10A). **(B)** Dot plots showing quantification of spine turnover (left), spine survival (middle) and spine density (right) under the four indicated treatments. Note that spines were imaged on the same dendrites across all conditions. Dots indicate mean ± SEM. Asterisks indicate significant differences to wakefulness in the left and middle panel. In the right panel, asterisks denote significant changes within each treatment compared to day 0. ^*^ p < 0.05, ^**^ p < 0.01,^***^ p < 0.001. **(C)** Imaging of acute spine dynamics during four different conditions. Left: schematic of the experimental timeline. Right: example of dendrite imaged during wakefulness in 10 min intervals (same dendrite as in A). **(D)** Dot plots showing quantification of acute spine turnover (left), spine survival (center) and spine density (right) under the four indicated treatments. Dots indicate mean ± SEM.

To rule out that the age of the animal influenced spine dynamics in the awake condition, we measured spine turnover in a group of age-matched animals to the first anesthesia group (Fig. S10A,C). Moreover, to rule out that the chronic imaging procedure per se and anesthesia in general had a long-lasting effect on the awake imaging condition, we added another awake-imaging control group with naïve, age-matched animals to the awake imaging time point in the experimental group (Fig. S10A,C). In all three groups, spine turnover was indistinguishable, indicating that neither age nor previous imaging under anesthesia impacted spine dynamics in the awake-imaging group (Fig. S10C).

Next, we asked whether the modulation of spine turnover by GAs was due to acute remodeling of spines during the time of anesthesia. Alternatively, spine turnover might be driven by long-lasting changes in network activity imposed by the slow reversal of all GAs. To capture fast events such as filopodia formation, we acquired image stacks every 10 min (Fig. 8C). Spine turnover, survival, or density were not significantly altered during the one hour of imaging (Fig. 8D). Thus, spines were stable during the one hour irrespective of the treatment. While mature spines typically show low elimination/formation rates over one hour, filopodia are more dynamic [64-66]. Unlike other reports, that observed an acute selective formation of filopodia under Keta/Xyl, but not Iso [67], we did not detect any acute effects of GAs on filopodia turnover of CA1 pyramidal cell dendrites. Thus, chronic exposure to all GAs consistently impacted spine dynamics, whereas acute effects were lacking. Keta/Xyl caused a strong decrease in spine turnover, accompanied by a higher surviving fraction and an increased density over time.

### Episodic memory consolidation is impaired by MMF and Keta/Xyl, but not by Iso

Episodic memory formation and consolidation require hippocampal activity. Newly learned experiences are thought of being consolidated via replay events that co-occur with low-frequency oscillations [49, 68-70]. In the hippocampus, these low-frequency events typically occur as sharp waves [50] during sleep, but also during awake resting behavior [70]. The above results from electrophysiological recordings and imaging showed that GAs strongly altered network oscillations in the CA1 area, in the case of MMF and Keta/Xyl, also long after anesthesia discontinuation. Spine turnover of CA1 pyramidal neurons was also affected, especially after Keta/Xyl administration. Therefore, we tested whether inducing anesthesia shortly after the acquisition of a new episodic memory affected its consolidation (Fig. 9A). In line with previous experiments, we restricted Iso and MMF anesthesia to one hour, while Keta/Xyl anesthesia was left to recede spontaneously. We assessed episodic-like memory with a water maze protocol for reversal learning, when the hidden platform was moved to the quadrant opposite the initial target location (Fig. 9A). Specifically, we tested the effects of the different anesthetics on the consolidation of the memory of the new platform location. We compared the performance of the mice during the probe trial done on day 3 immediately after the reversal learning protocol (and 30 min before anesthesia), with the performance during the probe trial on day 4, twenty-four-hours after anesthesia. During the probe trial on day 3, animals of all four groups spent significantly more time in the new target quadrant compared to chance (25 %), indicating that they learned the new platform position successfully (Fig. 9B,C).

**Fig 9.**
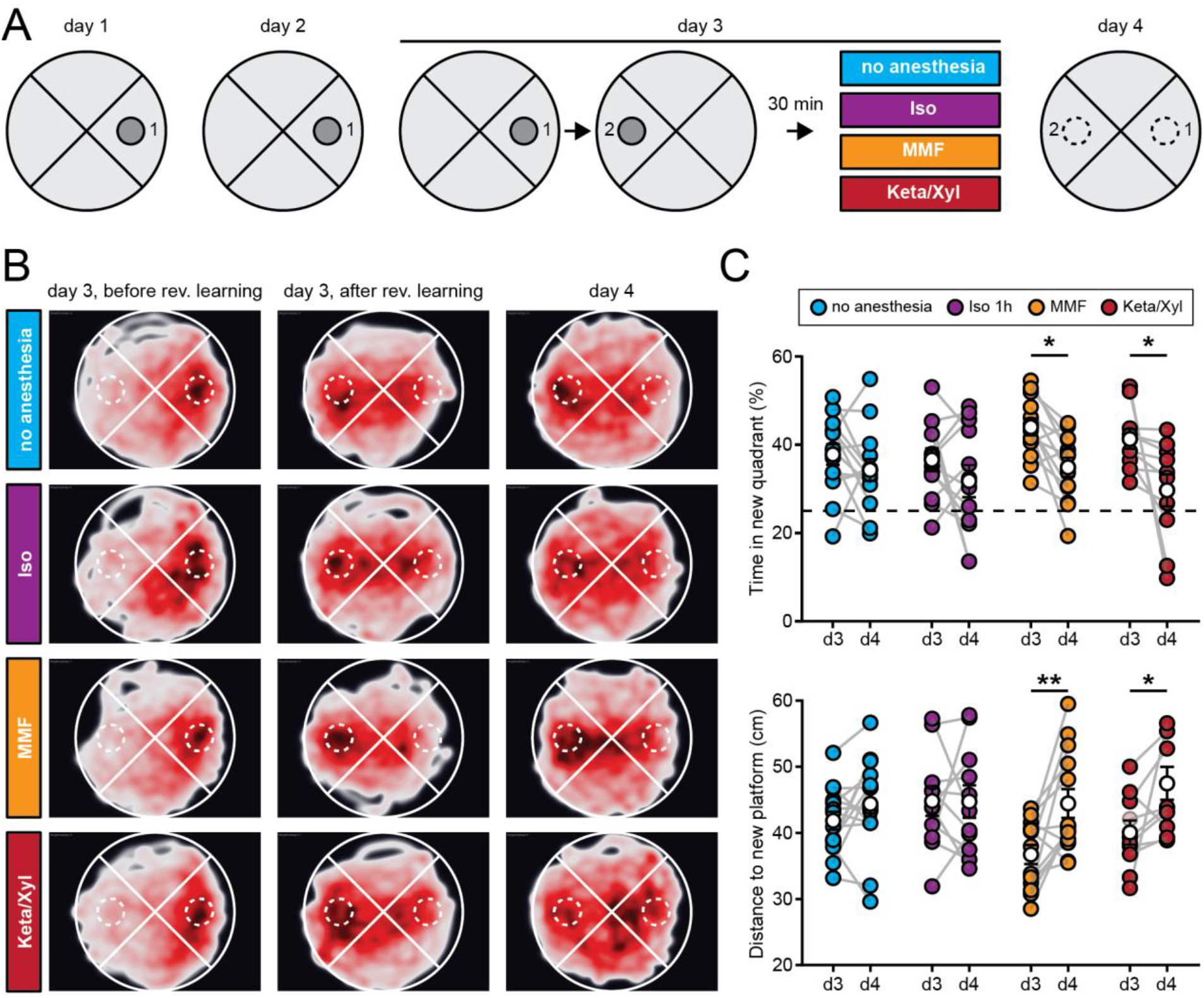
Episodic memory consolidation is impaired by MMF and Keta/Xyl, but not by Iso. **(A)** Experimental design to test episodic-like memory in a Morris water maze. On days 1 and 2, animals were trained to find the platform in position 1. Reversal learning was performed on day 3 where animals had to learn that the platform was moved to position 2. The training was followed 30 min later by a 1-h period of one of the four indicated treatments per group. On day 4, consolidation of the memory for the platform in position 2 was tested. **(B)** Heat maps showing trajectories of all mice during the first probe trial before reversal learning on day 3 (left column), after reversal learning on day 3 (middle column) and after treatment on day 4 (right column). The position of the target zone is indicated by dashed circles. **(C)** Scatter plots showing quantification of time spent in the new target quadrant (top) and distance to the new platform (bottom) after reversal learning on day 3 and on day 4. Filled, colored circles indicate individual animals, white circles indicate mean ± SEM. Asterisks in (C) indicate significant differences between days. ^*^ p < 0.05, ^**^ p < 0.01.

On day 4, control animals that did not undergo anesthesia showed the same performance as on day 3, suggesting that they had retained the memory of the new platform location (Fig. 9B,C). However, animals that were anesthetized with Keta/Xyl or MMF spent significantly less time in the new target quadrant and showed a significantly larger mean distance to the target platform position compared to the probe trial on day 3. In the Iso group, no significant difference compared to day 3 was detectable (Fig. 9B,C, S11A). The impairment of memory consolidation was not explained by the longer duration of recovery after Keta/Xyl or MMF compared to Iso, because anesthesia for up to 4 h with Iso had no disruptive effect (Fig. S11B,C). Thus, it is not the duration of the induced unconsciousness but rather the type of anesthetic that likely explains the impaired memory consolidation. Notably, the effects were relatively mild, and the decrease in performance on day 4 was not significantly different between treatment groups. In summary, consistent with long-lasting effects on CA1 network activity, Keta/Xyl, and MMF impaired episodic-like memory consolidation. In contrast, Iso, which overall caused a weaker disturbance of neuronal population activity and a faster recovery profile, did not significantly affect memory consolidation.

## DISCUSSION

We investigated and systematically compared the intra- and post-anesthetic effects of different commonly used anesthetic strategies on the mouse hippocampus across multiple levels of analysis. Despite sharing some common traits, brain and cellular network states differ substantially under the influence of various types of anesthetics [30, 71, 72]. Indeed, at the neuronal level, compared with awake state and natural sleep, all three anesthetics showed robustly reduced spiking activity in single neurons, reduced power in the high oscillation frequency band, and decorrelated cellular population activity. However, the induced network states in CA1 were highly distinct between the three different conditions, with Iso leading to prominent network oscillations at around 0.1 Hz, which timed the spiking activity of single units and neuronal calcium transients. Keta/Xyl caused pronounced oscillations between 0.5 and 4 Hz and the strongest reduction in calcium dynamics. MMF, in contrast, most strongly reduced LFP and SUA and impaired population dynamics for many hours as assessed with calcium imaging. Differences were also present in the long-term effects on spine dynamics, with Keta/Xyl stabilizing spines, leading to reduced turnover and increased density. MMF, on the other hand, mildly increased spine dynamics. Keta/Xyl cannot be antagonized and therefore changes of the CA1 network mediated by this anesthetic were present hours after the injection, in agreement with long-lasting overall changes of global animal physiology [38]. More unexpectedly, and in contrast to overall effects on physiology [38], CA1 network dynamics were still disturbed for at least 6 hours after antagonization of MMF anesthesia. These long-lasting alterations were associated with impairment of episodic memory consolidation after exposure to Keta/Xyl- or MMF, but not Iso. Thus, despite all fulfilling the same hallmarks of general anesthesia, different GAs distinctly alter hippocampal network dynamics, synaptic connectivity, and memory consolidation.

### Iso, MMF and Keta/Xyl have different molecular targets and distinctly modulate functional and structural features of CA1

The GAs used here represent three different strategies based on the large repertoire of currently available anesthetics. Isoflurane represents the class of halogenated diethyl ether analogues, which are volatile and therefore administered via inhalation. Fentanyl, in combination with the analgesic medetomidine and the sedative midazolam (MMF), represents an anesthetic approach based on the injection of a combination of drugs with sedative, analgesic and anxiolytic properties. In the clinic, propofol can be used instead of midazolam. Finally, ketamine is used both as an anesthetic and, at a lower dosage, as a treatment against depression. For anesthesia, it is generally combined with xylazine, which acts as a sedative, analgesic and muscle relaxant. All three strategies differ markedly in their molecular targets. Consequently, they uniquely modulate general animal physiology [38] and brain activity [72]. Isoflurane is a potent GABA- and glycine receptor agonist. Moreover, it activates two-pore potassium channels and acts as α-amino-3-hydroxy-5-methyl-4-isoxazolepropionic acid receptor (AMPAR) inhibitor [4]. Similar to Iso, midazolam, the hypnotic component of the MMF mix, mainly acts as a GABAR agonist with little effect on NMDARs. In contrast, ketamine is a potent, use-dependent NMDAR blocker with less pronounced effects on potassium channels, GABA, glycine and other glutamate receptors such as AMPA or kainite receptors [4]. Moreover, while most anesthetics reduce the activity of thalamic nuclei, ketamine increases thalamic drive [73], leading to enhanced rather than reduced oscillations in mid-to-high frequency bands such as theta and gamma [28, 74]. In accordance with this, our study reveals major differences in the action of the different anesthetics on functional and structural features of CA1. With both electrical recordings and calcium imaging we report a robust reduction of neuronal spiking and pairwise neuronal correlation. Notably, effects on electrical activity and calcium activity were well in line for both Iso and MMF, despite the different recording methods. However, we observed some divergence for Keta/Xyl.

### Comparison of electrophysiological recordings and calcium imaging

Generally, differences in electrophysiological recordings and calcium imaging data may stem from the location where the signal is detected. In the calcium imaging experiments, the signal was sampled in a horizontal plane located inside and parallel to stratum pyramidale of CA1. In this configuration, somatic, action-potential driven calcium transients mainly from pyramidal neurons dominate the signal. Due to the kinetics and calcium-binding properties of GCaMP6f, action potentials can only be resolved below approx. 5 Hz and are reported non-linearly [46, 75]. In contrast, the electrodes on linear probes are arranged orthogonally to the strata of CA1 and parallel to the dendrites of CA1 cells. Thus, synaptic potentials mainly constitute the LFP across all layers and spikes are picked up from both pyramidal cells (in stratum pyramidale) and GABAergic neurons in all layers. Moreover, the first method samples neurons that spatially distribute over a large area. In contrast, the second one is biased towards large, active neurons that are in close proximity of the electrode.

More specifically, under Keta/Xyl, the overall firing rate of single units showed the smallest reduction of all three anesthetics. At the same time, imaging revealed the most substantial reduction in rate, amplitude and duration of calcium transients (compare Fig. 2B and 3D). One reason for this discrepancy may be the inhibitory action of ketamine on NMDARs. CA1 pyramidal cells display large, NMDAR-driven dendritic plateau potentials and calcium spikes [76]. Moreover, ketamine likely inhibits L-type voltage-gated calcium channels [77] and reduces burst firing [78], leading to calcium transients with reduced amplitude and a faster decay constant. In contrast, ketamine has little influence on sodium spikes and AMPAR-mediated synaptic potentials, which are detected in electrical recordings as SUA and LFP, respectively. In accordance with electrical recordings, calcium transients showed increased power at 0.1-0.2 Hz under Iso. However, we did not detect a clear peak at 1-4 Hz in the presence of Keta/Xyl, as seen in LFP and SUA, probably due to its strongly dampening effect on calcium transients. The (low-pass) filtering of neuronal activity imposed by calcium indicators might also play a role [75].

Notably, the differences between electrical recordings and calcium imaging under Keta/Xyl are relevant. Calcium is a second messenger central to neuronal plasticity and metabolism [79, 80]. NMDARs are a major source for activity-dependent calcium entry into the cell, involved in regulating synaptic plasticity, metabolism, and pathology [81]. The present findings suggest that Keta/Xyl has a particularly strong effect on neuronal calcium activity, uncoupling action potential firing from associated cytosolic calcium transients, leading to reduced intracellular calcium signaling. In contrast, calcium transients under MMF and Iso anesthesia closely matched the electrical activity profile of neurons. Therefore, aside from overall effects on network activity, Keta/Xyl may selectively alter neuronal plasticity by suppressing NMDAR-dependent postsynaptic calcium signals.

### In contrast to neocortex, GAs decorrelate neuronal activity in CA1

All anesthetics decorrelated neuronal activity in CA1, leading to an overall fragmented network state with an increased number of temporal and spatial clusters. This is in stark contrast with what has been reported from studies on GAs and cortical activity both at adulthood [25-27] and during development [58]. This discrepancy may arise from the distinct architecture of CA1 compared to L2/3 of the neocortex, the latter showing a high degree of local interconnectivity [47]. In CA1 this is not the case. Pyramidal cells receive their main excitatory input from CA3 and entorhinal cortex and send their efferents to subiculum and extrahippocampal areas without making local connections among each other [9]. Afferent activity originating in various sources and converging in CA1, may arrive out-of-phase under anesthesia, leading to desynchronized firing of CA1 pyramidal cells. Such a phenomenon has been proposed as a candidate mechanism underlying desynchronization of neuronal firing in basal ganglia under conditions of slow oscillations (slow-wave sleep) and high synchrony in the neocortex [82].Notably, the pairwise correlation was not entirely independent of the distance between neurons. Synchronization of pyramidal neurons via local, GABAergic interneurons may be another factor that increases spatial correlations. Both in the neocortex and hippocampus, various types of GABAergic interneurons locally connect to and synchronize pyramidal neurons such as basket or bistratified cells [83].

Coordinated neuronal network dynamics, including pairwise correlation of calcium transients and single units, population coupling, clustering in the temporal and spatial domain were consistently impaired most strongly with Keta/Xyl and MMF. Iso, both in electrophysiological as well as calcium recordings, showed the mildest effects and permitted hippocampal activity patterns that most closely resembled wakefulness and NREM/REM sleep. Iso and MMF, in contrast to Keta/Xyl, are thought to be immediately reversible [38]. However, especially MMF showed significant disruption of network dynamics long after reversal both in electrical recordings and with calcium imaging. Antagonization of MMF failed to fully recover calcium dynamics within the following 5 hours. Such long-lasting alterations might interfere with the hippocampal function shortly after antagonization of MMF and must be considered when performing whole-cell recordings in freely moving animals [84-86].

Since all anesthetics had a much longer effect on network activity than we expected, we asked whether this is reflected in long-term effects of these different types of anesthetics on spine dynamics of CA1 pyramidal neurons. Recent studies investigating spine dynamics at CA1 pyramidal neurons came to incongruent conclusions reporting spine turnover ranging from 3% [63] over 12% [21] to approx. 80% [62] over 4 days. However, all studies used either isoflurane [21] or ketamine/xylazine-based [62, 63] anesthesia during the repeated imaging sessions. Thus, to what extent anesthesia itself influences spine dynamics is not clear.

### Iso, MMF and Keta/Xyl distinctly alter spine dynamics in CA1

More generally, various effects of general anesthesia on spine dynamics were reported, depending on the brain region, preparation, age of the animal and anesthetic strategy. For example, enhanced synaptogenesis has been reported with different types of anesthetics on cortical and hippocampal neurons during development [59, 60]. In contrast, one study indicated that spine dynamics were not altered on cortical neurons of adult mice with Keta/Xyl or Iso [67], while another study demonstrated an increase in spine density in somatosensory cortex with ketamine [87]. Also, fentanyl-mediated, concentration-dependent bidirectional modulations of spine dynamics were reported in hippocampal cultures [88].

To systematically compare spine dynamics in CA1 in vivo under different anesthetic treatments, we imaged spines at basal, oblique and tuft dendrites in a large set of dendrites. We found small, but robust chronic effects of repeated anesthesia. These alterations were present in all strata of CA1, consistent with a layer-independent reduction of SUA during anesthesia. Keta/Xyl decreased spine turnover leading to a mild increase in spine density over time by stabilizing existing spines. This observation agrees with recent studies that showed a stabilizing effect of ketamine in the somatosensory cortex, resulting in increased spine density [87]. Thus, repeated anesthetic doses of Keta/Xyl may limit overall synaptic plasticity and thus spine turnover. It was further shown that sub-anesthetic, antidepressant doses of ketamine enhance spine density in the prefrontal cortex [89, 90], similar to our study of CA1 neurons. Iso and MMF had contrasting effects on spine dynamics compared to Keta/Xyl, mildly enhancing spine turnover, which might be explained by their different pharmacology compared to ketamine, as pointed out above. A second aspect that distinguishes Keta/Xyl from Iso and MMF is its irreversibility, which might lead to longer-lasting alterations of synaptic transmission and E/I ratios leading to differential spine dynamics. This idea is supported by the observation that during the anesthesia period itself, spine turnover was not altered, suggesting that long-lasting and repeated disturbances are required to leave a mark in synaptic connectivity.

### MMF and Keta/Xyl, but not Iso, retrogradely affect episodic-like memory formation

Sleep is a natural form of unconsciousness and is required for memory consolidation, including hippocampus-dependent memories [49, 56]. Recent work suggested that sleep- and anesthesia-promoting circuits differ [91, 92] while others identified circuit elements shared between sleep and general anesthesia [93], especially during development [58]. Therefore, we asked how the diverse alterations of CA1 network dynamics imposed by the different anesthetics impact memory consolidation. In our study Iso resembled most closely network states during wakefulness and natural sleep, while Keta/Xyl and MMF caused strong, lasting alterations of LFP, SUA and calcium dynamics.

Notably, a single dose of anesthesia with Keta/Xyl and MMF, but not Iso disrupted memory consolidation using a water maze assay in adult mice. Retrograde amnesia appeared to be more sensitive to the magnitude than the duration of CA1 network disturbance imposed by the various anesthetics. Keta/Xyl and MMF most strongly decorrelated CA1 network activity and reverted only slowly. Extending the duration of Iso anesthesia up to 4 h, to match the slow recovery after MMF and Keta/Xyl, did not affect memory consolidation. This observation indicates that the slow recovery of network activity after Keta/Xyl and MMF alone cannot explain anesthesia-mediated disruptions of memory consolidation. Instead, specific aspects of the different anesthetics may selectively impact hippocampus-dependent memory formation. For example, ketamine is an NMDAR blocker that has been shown to be necessary for the long-term stabilization of place fields in CA1 [94], encoding of temporal information of episodes [95], and formation of episodic-like memory [96].

Our results appear at odds with a report [97], where a single, 1-h treatment with Iso caused deficits in the formation of contextual fear memory, object recognition memory and performance in the Morris water maze in the following 48 h. However, this study investigated memory acquisition after anesthesia (i.e., anterograde amnesia), while our study asked whether anesthesia affects the consolidation of a memory formed shortly before the treatment (i.e., retrograde amnesia).

Changes in synaptic connections are considered essential for memory formation and storage [11-14]. Despite a small effect on spine dynamics, the strong and lasting disturbance of hippocampal network activity in CA1 (and most likely other brain areas) by Keta/Xyl and MMF was sufficient to interfere with memory consolidation. The chronic alterations of spine turnover, especially by Keta/Xyl, may therefore indicate that repeated anesthesia can impact long-lasting hippocampus-dependent memories.

To establish a direct link between spine dynamics, network disruptions and memory, future studies are required that investigate both spine turnover and changes in population coupling at hippocampal neurons causally involved in memory formation and maintenance.

Taken together, we report a novel effect of anesthesia on brain dynamics, namely fragmentation of network activity in hippocampus. We consistently observe this phenomenon across multiple levels of analysis. This unique response compared to the cortex may underlie its high sensitivity to anesthesia, including its central role in amnesia. The extent, duration, and reversibility of network fragmentation depend on the GA used. Therefore, this study may help guide the choice of an appropriate anesthetic strategy, dependent on experimental requirements and constraints, especially in the neurosciences. More generally, our findings might also have relevance for the clinic. Postoperative delirium, a condition that involves memory loss, is still an unresolved mystery. Minimizing the disturbance of hippocampal function may be one building block to overcome this undesired condition.

## AUTHOR CONTRIBUTIONS

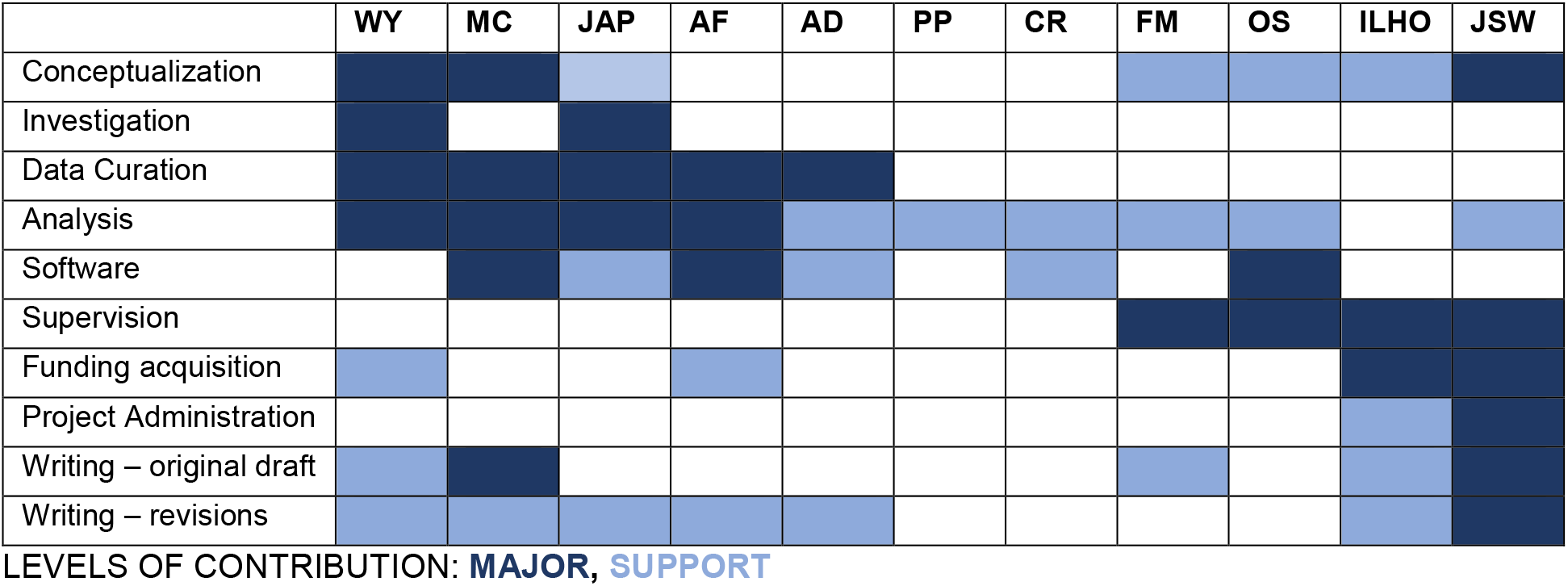

## ACKNOWLEDGEMENTS

We thank Stefan Schillemeit and Kathrin Sauter for technical assistance and Thomas G. Oertner and Amit Marmelstein for critical feedback on the manuscript. This work was funded by the Deutsche Forschungsgemeinschaft (DFG, SPP1926, FOR2419/P6, SFB963/B8 to J.S.W., SPP 1665/Ha 4466/10-1/Ha4466/12-1, SFB 936/B5 to I.L.H.-O., SFB 936/B7 to F.M.), the European Research Council (ERC2016-StG-714762 to J.S.W., ERC-2015-CoG 681577 to I.L.H.-O.), the German Academic Exchange Service (DAAD, STG/19/5744091 to A.F.), and the Chinese Scholarship Council (CSC 201606210129 to W.Y.).

## METHODS

### Experimental Models and Methods

### Mice

Adult C57BL/6J mice and transgenic Thy1-GFP-M mice of both sexes were housed and bred in pathogen-free conditions at the University Medical Center Hamburg-Eppendorf. The light/dark cycle was 12/12 h and the humidity and temperature were kept constant (40% relative humidity; 22°C). Food and water were available ad libitum. All procedures were performed in compliance with German law according and the guidelines of Directive 2010/63/EU. Protocols were approved by the Behörde für Gesundheit und Verbraucherschutz of the City of Hamburg.

### Hippocampal recording-window surgery and in-vivo electrophysiology

Chronic multisite extracellular recordings were performed in dorsal CA1 during the dark phase of the dark/light cycle, except for sleep recordings, which were done during the light period. The adapter for head fixation was implanted at least 4 days before recordings. Mice were anesthetized via intraperitoneal injection of midazolam/medetomidine/fentanyl (MMF) and placed on a heating blanket to maintain the body temperature. Eyes were covered with eye ointment (Vidisic, Bausch + Lomb) to prevent drying. Prior to surgery, the depth of anesthesia and analgesia was evaluated with a toe-pinch to test the paw-withdrawal reflex. Subsequently, mice were fixed in a stereotactic frame, the fur was removed with a fine trimmer and the skin of the head was disinfected with Betaisodona. After removing the skin, 0.5% bupivacaine / 1% lidocaine was locally applied to cutting edges. A metal head-post (Neurotar) was attached to the skull with dental cement (Super Bond C&B, Sun Medical) and a craniotomy was performed above the to the dorsal CA1 area (−2.0 mm AP, ± 1.3 mm ML relative to Bregma) which was subsequently protected by a customized synthetic window filled with Kwik-Cast sealant (World Precision Instruments). After recovery from anesthesia, mice were returned to their home cage and were provided with Meloxicam mixed into soft food for 3 days. After recovery from the surgery, mice were accustomed to head-fixation and trained to move in the Mobile HomeCage system (Neurotar). For recordings, craniotomies were reopened by removal of the Kwik-Cast sealant and multi-site electrodes (NeuroNexus, MI, USA) were inserted into the dorsal CA1 (one-shank, A1×16 recording sites, 50 µm spacing, 1.6 mm deep). A silver wire served as ground and reference in the craniotomy between skull and brain tissue. Extracellular signals were band-pass filtered (0.1-8000 Hz) and digitized (32 kHz) with a multichannel extracellular amplifier (Digital Lynx SX; Neuralynx). The same animals were recorded weekly under different anesthesia. After 15 min of non-anesthetized recording, mice received a subcutaneous injection of Keta/Xyl, MMF or inhalation of Iso in a pseudo-randomized order. The following drug combinations were administered: 2.0 % isoflurane in 100% O_2_; 130 mg/kg ketamine, 10 mg/kg xylazine s.c.; 5.0 mg/kg midazolam, 0.2 mg/kg medetomidine and 0.05 mg/kg fentanyl s.c.; and for complete reversal of anesthesia, 0.5 mg/kg flumazenil, 2.5 mg/kg atipamezole and 0.1 mg/kg buprenorphine s.c. Recordings were conducted for 1.5 h. After recordings, the craniotomy was closed and mice were returned to their home cage. Electrode position was confirmed in brain slices postmortem.

### EMG recordings

Electromyography (EMG) electrodes for sleep state classification were implanted during hippocampal recording-window surgery. Two gold plates (∼3 mm diameter) soldered to epoxy lacquered wires and attached to a connector were inserted into the right and left nuchal muscles and fixed with dental cement (Super Bond C&B, Sun Medical). For EMG recordings, a cable was attached to the implanted connector and directly digitized (32 kHz) and band-pass filtered (8-8000 Hz) through a customized break-out channel board with a multichannel amplifier (Digital Lynx SX; Neuralynx). EMG recordings were done at least for 1 h 45 min in non-anesthetized mice without disturbances. Mice were recorded 1-2 times.

### Virus injection and hippocampal window surgery for in vivo calcium imaging

C57BL/6J wild-type mice were anesthetized via intraperitoneal injection of midazolam/medetomidine/fentanyl (MMF) and placed on a heating blanket to maintain the body temperature. Eyes were covered with eye ointment (Vidisic, Bausch + Lomb) to prevent drying. Prior to surgery, the depth of anesthesia and analgesia was evaluated with a toe-pinch to test the paw-withdrawal reflex. Subsequently, mice were fixed in a stereotactic frame, the fur was removed with a fine trimmer and the skin of the head was disinfected with Betaisodona. The skin was removed by a midline scalp incision (1-3 cm), the skull was cleaned using a bone scraper (Fine Science Tools) and a small hole was drilled with a dental drill (Foredom) above the injection site. AAV2/7-syn-GCaMP6f was targeted unilaterally to the dorsal CA1 area (- 2.0 mm AP, ± 1.3 mm ML, - 1.5 mm DV relative to Bregma). 0.6 µl of virus suspension was injected. All injections were done at 100 nl^*^min^-1^ using a glass micropipette. After the injection, the pipette stayed in place for at least 5 min before it was withdrawn and the scalp was closed with sutures. For complete reversal of anesthesia, mice received a subcutaneous dose of Flumazenil, Atipamezole and Buprenorphine (FAB). During the two days following surgery animals were provided with Meloxicam mixed into soft food. Two weeks after virus injection, mice were anesthetized as described above to implant the hippocampal window. After fur removal, skin above the frontal and parietal bones of the skull was removed by one horizontal cut along basis of skull and two rostral cuts. The skull was cleaned after removal of the periosteum, roughened with a bone scraper and covered with a thin layer of cyanoacrylate glue (Pattex). After polymerization a 3-mm circle was marked on the right parietal bone (anteroposterior, −2.2 mm; mediolateral, +1.8 mm relative to bregma) with a biopsy punch and the bone was removed with a dental drill (Foredom). The dura and somatosensory cortex above the hippocampus were carefully aspirated until the white matter tracts of the corpus callosum became visible. The craniotomy was washed with sterile PBS and a custom-built imaging window was inserted over the dorsal hippocampus. The window consisted of a hollow glass cylinder (diameter: 3 mm, wall thickness: 0.1 mm, height: 1.8 mm) glued to a No. 1 coverslip (diameter: 3mm, thickness: 0.17 mm) on the bottom and to a stainless-steel rim on the top with UV-curable glass glue (Norland NOA61). The steel rim and a head holder plate (Luigs & Neumann) were fixed to the skull with cyanoacrylate gel (Pattex). After polymerization, cranial window and head holder plate were covered with dental cement (Super Bond C&B, Sun Medical) to provide strong bonding to the skull bone. Following the surgery, animals were provided with Meloxicam mixed into soft food for 3 days. The position of the hippocampal window was confirmed in brain slices postmortem.

### Two-photon calcium imaging in anesthetized and awake mice

The same animals were sequentially imaged under Keta/Xyl, MMF or Iso in a pseudo-randomized order during the dark phase of the dark/light cycle (for details, see above). After losing the righting reflex, generally 5–10 min after application of the anesthetics, the animals were positioned on a heating-pad to maintain body temperature at approximately 37°C during anesthesia. The intensity of anesthesia and evaluation of the different stages of anesthesia were assessed by recording the presence or absence of distinct reflex responses: righting reflex, palpebral reflex, toe-pinch reflex. Between each imaging session, mice were allowed to recover for one week.

Anesthetized mice were head-fixed under the microscope on a heated blanket to maintain body temperature. Eyes were covered with eye ointment (Vidisic, Bausch + Lomb) to prevent drying. The window was centered under the two-photon microscope (MOM-scope, Sutter Instruments, modified by Rapp Optoelectronics) and GCaMP6f expression was verified in the hippocampus using epi fluorescence. Images were acquired with a 16x water immersion objective (Nikon CFI75 LWD 16X W, 0.80 NA, 3.0 mm WD). For awake imaging we used a linear treadmill, which allowed imaging during quiet and running states. 5-min-timelapse images were acquired every 10 minutes for a period of 50 minutes. Only quiet periods were considered for analysis in this study. Image acquisition was carried out with a Ti:Sa laser (Chameleon Vision-S, Coherent) tuned to 980 nm to excite GCaMP6f. Single planes (512×512 pixels) were acquired at 30 Hz with a resonant-galvanometric scanner at 29-60 mW (980 nm) using ScanImage 2017b (Vidrio). Emitted photons were detected by a pair of photomultiplier tubes (H7422P-40, Hamamatsu). A 560 DXCR dichroic mirror and a 525/50 emission filter (Chroma Technology) was used to detect green fluorescence. Excitation light was blocked by short-pass filters (ET700SP-2P, Chroma). For the repetitive imaging, the position of the field of view (FOV) was registered in the first imaging session with the help of vascular landmarks and cell bodies of CA1 pyramidal neurons. This allowed for subsequent retrieval of the FOV for each mouse.

Calcium imaging experiments to measure recovery from anesthesia were done in five additional animals. They were trained to maintain immobile on the treadmill for extended periods. We ensured to measure the same FOV and to maintain overall stability of fluorescence intensity for every recording in each imaging session for a given animal. The time-lapse recordings were extended to up to a maximum of 10 min per time point to have a higher probability of capturing motionless periods continuously, in awake and recovery states. Iso was applied for 60 min. FAB was injected 60 min after the application of MMF. Keta/Xyl was not antagonized. Imaging of calcium activity was performed before, 0.5, 1.5, 2, 3, 4, 5, and 6 hours after induction of anesthesia. For Iso and MMF, the 1.5 h time point represented the first imaging session after reversal. Untreated control animals were imaged every hour for the same amount of time.

To habituate mice to sleep under head-fixation, we used a linear treadmill, which allowed the mice to move at will. Through the first 4 sessions mice were kept head-fixed for 15 to 30 min. In ten following sessions the fixation period was extended up to 4h with increasing intervals of 30 min. The state of the mouse was continuously monitored with a USB camera and the running speed was recorded with custom-written scripts in the Matlab. After habituation to 4h head-fixation, sleep imaging sessions were recorded, which were synchronized with recordings of the pupil and running speed. Sleep imaging was performed during the light phase of the dark/light cycle.

### Two-photon spine imaging in anesthetized and awake mice

3 - 4 weeks after window implantation, chronic spine imaging started in Tg(Thy1-EGFP)MJrs/J mice with the first of a total of four imaging series (see Fig. S10A). Each imaging series was done under one of the three anesthetic conditions (Iso, Keta/Xyl, MMF, see above for details) or during wakefulness. Within one series, mice were imaged 5 times every 4 days. Afterwards, mice were allowed to recover for three to four weeks until the next imaging series under a different anesthetic condition was started. Thus, each experiment lasted approx. 5 months. To avoid time-dependent effects, anesthetic conditions were pseudo-randomized (see Fig. S10A). For imaging sessions under anesthesia mice were head fixed under the microscope on a heated blanket to maintain body temperature. Eyes were covered with eye ointment (Vidisic, Bausch + Lomb) to prevent drying. The window was centered under the two-photon microscope (MOM-scope, Sutter Instruments, modified by Rapp Optoelectronics) and GFP expression was verified in the hippocampus using epi-fluorescence. Image acquisition was carried out with a Ti:Sa laser (Chameleon Vision-S, Coherent) tuned to 980 nm to excite GFP. Images were acquired with a 40x water immersion objective (Nikon CFI APO NIR 40X W, 0.80 NA, 3.5 mm WD). Single planes (512×512 pixels) were acquired at 30 Hz with a resonant scanner at 10-60 mW (980 nm) using ScanImage 2017b. Before the first imaging session, we registered the field of views with the help of vascular landmarks and cell bodies of CA1 pyramidal neurons and selected several regions for longitudinal monitoring across the duration of the time-lapse experiment. Each of these regions contained between 1 and 2 dendritic segments visibly expressing GFP. The imaging sessions lasted for max 60 min and mice were placed back to their home cages where they woke up.

### Morris Water Maze

We designed a protocol for reversal learning in the spatial version of the water maze to assess the possible effects of the different anesthetics on episodic-like memory in mice [98, 99]. The water maze consisted of a circular tank (145 cm in diameter) circled by dark curtains and walls. The water was made opaque by the addition of non-toxic white paint such that the white platform (14 cm diameter, 9 cm high, 1 cm below the water surface) was not visible. Four landmarks (35 × 35 cm) differing in shape and grey gradient were positioned on the wall of the maze. Four white spotlights on the floor around the swimming pool provided homogeneous indirect illumination of 60 lux on the water surface. Mice were first familiarized for one day to swim and climb onto a platform (diameter of 10 cm) placed in a small rectangular maze (42.5 × 26.5 cm and 15.5 cm high). During familiarization, the position of the platform was unpredictable since its location was randomized, and training was performed in darkness. After familiarization, mice underwent three learning days, during which they had to learn the location of a hidden platform. The starting position and the side of the maze from which mice were taken out of the maze were randomized. On day 1, mice underwent four learning trials (maximum duration of 90 seconds, inter-trial interval of 10 minutes). After staying on the platform for 15 s, mice were returned to their home cage and warmed up under red light. On day 2, mice underwent two training trials before they performed a 60 seconds-long probe trial to assess their searching strategy. Afterwards, one additional training trial was used to re-consolidate the memory of the platform position, and mice were distributed into four groups with a similar distribution of performance. On day 3, the long-term memory of the platform position was tested with a 45-seconds long probe trial, followed by another training trial with the platform in place to avoid extinction. Then mice underwent four reversal learning trials with the platform located in the quadrant opposite the one in which the platform was during the previous training trials. To assess whether the mice learned the new platform position, mice underwent a 60-seconds long probe trial followed by one more training trial to consolidate the memory of the new location. One hour after the last reversal learning trial, mice were anesthetized to analyze the effects of the anesthesia on the consolidation of the memory of the new platform position. Mice were assigned to four groups with an equal average performance during the probe trial on day 2. Each group was subjected to different conditions: one-hour Iso anesthesia, one-hour MMF anesthesia, Keta/Xyl anesthesia (which was not antagonized), and one group was left untreated. On day 4, mice underwent a 60-seconds long probe trial to evaluate their searching strategies; namely, the “episodic-like memory” of the reversal learning trials performed one hour before having been anesthetized on day 3 (see Fig. 9A).

### Quantification and Statistical Analysis

#### Electrophysiology

In vivo electrophysiology data were analyzed with custom-written scripts in the Matlab environment available at https://github.com/mchini/HanganuOpatzToolbox. We selected the recording site in the pyramidal layer of CA1. Data were band-pass filtered (1-100 Hz or 0-100 Hz for low frequency LFP analysis) using a third-order Butterworth forward and backward filter to preserve phase information before down-sampling to analyze LFP.

##### Detection of active periods

Active periods were detected with an adapted version of an algorithm for ripple detection (https://github.com/buzsakilab/buzcode/blob/master/detectors/detectEvents/bz_FindRipples.m). Briefly, active periods were detected on the band-pass filtered (4-20 Hz) normalized squared signal using both absolute and relative thresholds. We first passed the signal through a boxcar filter and then performed hysteresis thresholding: we first detected events whose absolute or relative power exceeded the higher threshold, and considered as belonging to the same event all data points that were below the lower (absolute or relative) threshold. Absolute thresholds were set to 7 and 15 μV, relative thresholds to 1 and 2. Periods were merged if having an inter-period interval shorter than 900 ms, and discarded if lasted less than 500 ms. Percentage of active periods was calculated for 15 min bins. Timestamps were preserved for further analysis.

##### Power spectral density

Power spectral density was calculated on 30 s-long windows of 0-100 Hz filtered signal using Welch’s method with a signal overlap of 15 s.

##### Modulation index (MI)

Modulation index was calculated as (value anesthesia - value pre-anesthetized) / (value anesthesia + value pre-anesthetized).

##### Power law decay exponent of the LFP power spectrum

The 1/f slope was computed as in [36]. We used robust linear regression (Matlab function *robustfit*.*m*) on the log10 of the LFP power spectrum in the 30-50 Hz frequency range.

##### Phase-amplitude coupling (PAC)

PAC was calculated on 0-100 Hz filtered full signal using the PAC toolbox based on modulation index measure [100]. Range of phase vector was set to 0-8 Hz and range of amplitude vector was set to 20-100 Hz. Significant coupling was calculated in comparison to a shuffled dataset. Non-significant values were rejected.

##### Single unit analysis

Single unit activity (SUA) was detected and clustered using klusta [101] and manually curated using phy (https://github.com/cortex-lab).

##### Active units

the recording was divided into 15-minute bins. Single units were considered to be active in the time interval if they fired at least five times.

##### Pairwise phase consistency

Pairwise phase consistency (PPC) was computed as previously described [102]. Briefly, the phase in the band of interest was extracted as mentioned above, and the mean of the cosine of the absolute angular distance (dot product) among all single unit pairs of phases was calculated.

##### Unit Power

SUA spike trains of each recording were summed in a population vector, and power spectral density was calculated on 30 s-long windows using Welch’s method with a signal overlap of 15 s. The resulting power spectra were normalized by the firing rate in that window.

##### Spike-Time tiling coefficient

(STTC) was computed as previously described [52]. Briefly, we quantified the proportion (P_A_) of spikes of spike train A that fall within ±Δt of a spike from spike train B. To this value we subtract the proportion of time that occurs within ±Δt of spikes from spike train B (T_B_). This is then divided by 1 minus the product of these two values. The same is then applied after inverting spike train A and B, and the mean between the two values is kept.

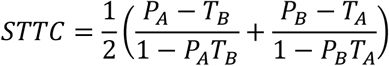

Importantly, this coefficient has several desirable properties. It is bounded between −1 and 1. It is symmetric with respect to the two spike trains. Computing it over different timescales is readily done by controlling the value of the parameter “Δt”. Lastly, and most importantly, traditionally used methods of assessing correlations between pairs of spike trains show an inverse correlation between their value and firing rate, due to the fact that spiking is sparse with respect to the sampling frequency, and therefore quiescent period in both spike trains artificially increase the correlation. This is not the case for the spike-time tiling coefficient [52]. Given that there are large differences in the average firing rate of our conditions, we chose STTC analysis over pure correlation analysis to circumvent this major bias. On the flipside, STTC cannot be straightforwardly applied to negative correlations, that were therefore not investigated in SUA data.

#### Calcium imaging data

In vivo calcium imaging data were analyzed with custom-written scripts in the Python and Matlab environment available at https://github.com/mchini/Yang_Chini_et_al.

##### Alignment of multiple recordings

To track the activity of the same set of neurons in different anesthetic conditions and during wakefulness, we acquired two-photon time series of a defined field of view for each animal and each condition across multiple weeks. Over such long time periods, the field of view was susceptible to geometrical transformations from one recording to another and thus, any two time series were never perfectly aligned. This problem scaled with time that passed between recordings. However, optimal image alignment is critical for the successful identification and calcium analysis of the same neurons across time [103, 104].

To address this problem, we developed an approach based on the pystackreg package, a Python implementation of the ImageJ extension TurboReg/StackReg [105]. The source code that reproduces the procedure described in this section is available on github (https://pypi.org/project/pystackreg/). The *pystackreg* package is capable of using different combinations of geometrical transformations for the alignment. We considered rigid body (translation + rotation + scaling) and affine (translation + rotation + scaling + shearing) transformation methods, which we applied to mean and enhanced-mean intensity images generated by Suite2p during the registration of each single recording. We performed the alignment using all four combinations (2 transformations x 2 types of images) choosing the one with the best performance according to the following procedure. Squared difference between the central part of a reference and aligned image served as a distance function d to quantify the alignment (since the signal is not always present on the borders of the image they were truncated):

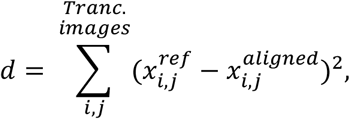

where 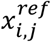 and 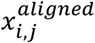 are intensities of the pixel with coordinates *i j* of the reference and aligned images. The combination with the smallest score was chosen for the final transformation. In some rare cases, the algorithm of the alignment did not converge for a given transformation method and image type (mean or enhanced-mean), crumbling the aligned image in a way that most of the field of view remained empty. This combination may have the smallest distance function *d* and may be falsely identified as the best one. To overcome this issue, an additional criterion was applied, which requires the central part of the aligned picture to contain more than 90 % of the non-empty pixels. The overall performance of the algorithm was verified by visual inspection. An example of the alignment of two recordings is shown in Fig. S5. The alignment for all recordings of an example mouse is demonstrated in a supplementary video (Supplementary_video_37529_aligned_recordings.avi).

In case of relatively small distortions across recordings, for example, when consecutive acquisitions were done within one imaging session, registration can alternatively be performed simultaneously with ROI detection in Suite2p by concatenating those TIFF-stacks. In this approach, every ROI is automatically labeled with the same identification number across all recordings.

##### Identification of the same neurons across different recordings & unique neuron ID assignment

After the alignment procedure, we set out to identify neurons which were active across multiple recordings (and thus, multiple conditions). To achieve this, we developed an algorithm similar to the one described in Sheintuch. et al. 2017 [104]

The algorithm processes in series all recordings for a given animal and assigns unique identification (ID) numbers to each ROIs of every recording. Since the recordings under Iso-anesthesia had the largest number of active neurons, we chose the first recording of this condition as reference. We assigned IDs that ranged from 1 to the total amount of neurons to all the ROIs of this recording. For every other recording of each mouse, Neuron ID assignment consisted of: 1. comparison of the properties (details below) of each ROI with each ROI that had already been processed. 2a. If the properties of the ROI matched the properties of an ROI from a previously analyzed recording, the ROI received the same Neuron ID. 2b. If no match was found, a new (in sequential order) Neuron ID was assigned to the ROI. In order to be identified as representing the same neuron in two different recordings, two ROIs had to respect the following criteria: the distance between their centroids had to be below 3 µm, and the overlap between their pixels had to be above 70%. An example of the identification of unique neuron pairs in two recordings is presented in Fig. S6A. The thresholds were chosen based on the distribution of the distances between centroids and percentage of the overlaps. An example for a single mouse is graphically illustrated in Fig. S6B. Both properties have a clearly bimodal distribution (similar to [104]) with cutoffs close to the chosen thresholds.

##### Signal extraction and analysis

Signal extraction, correlation and spectral analysis for calcium signal was performed using Python (Python Software Foundation, NH, USA) in the Spyder (Pierre Raybaut, The Spyder Development Team) development environment. Calcium imaging data were analyzed with the Suite2p toolbox [106] using the parameters given in table 1.

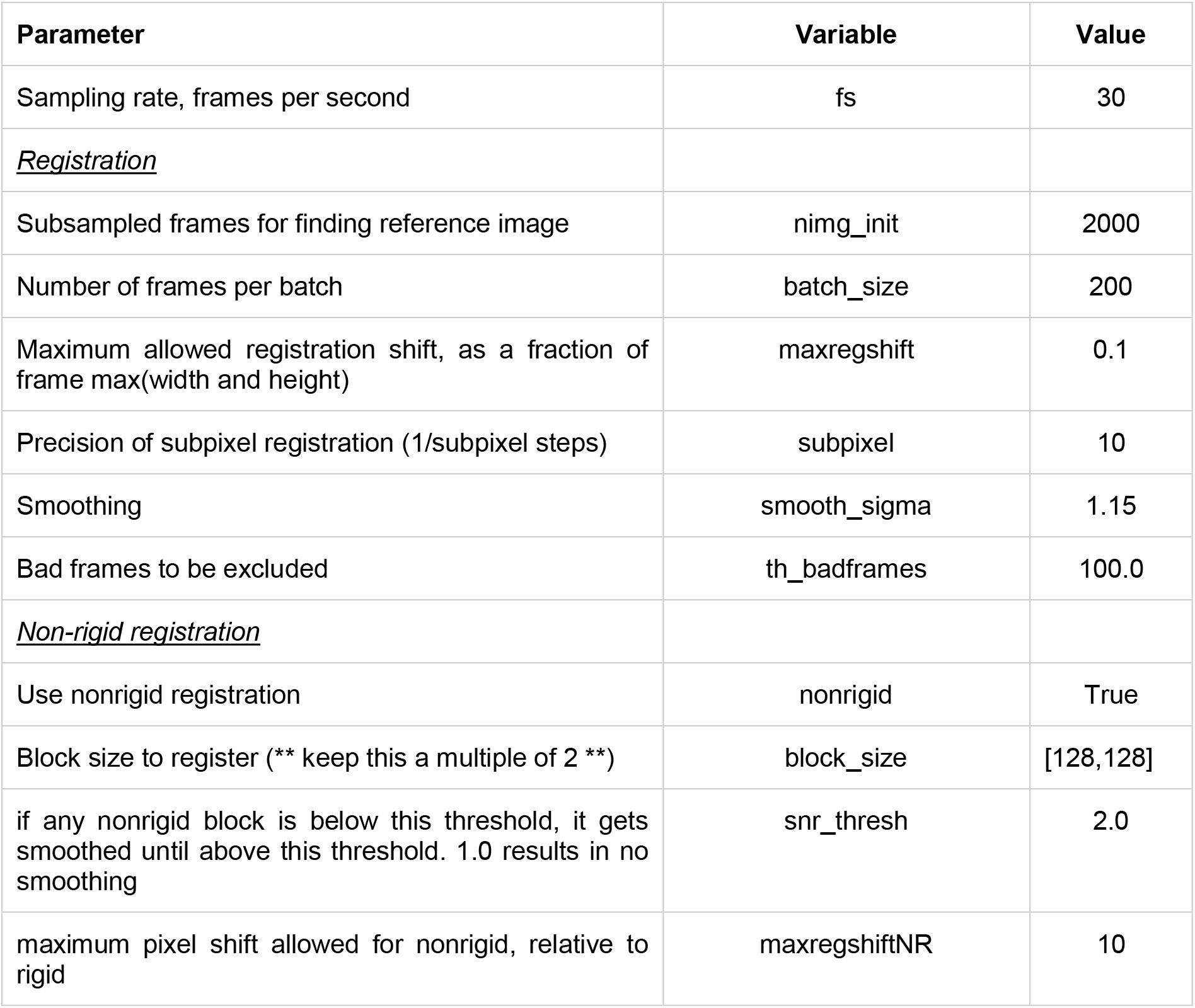
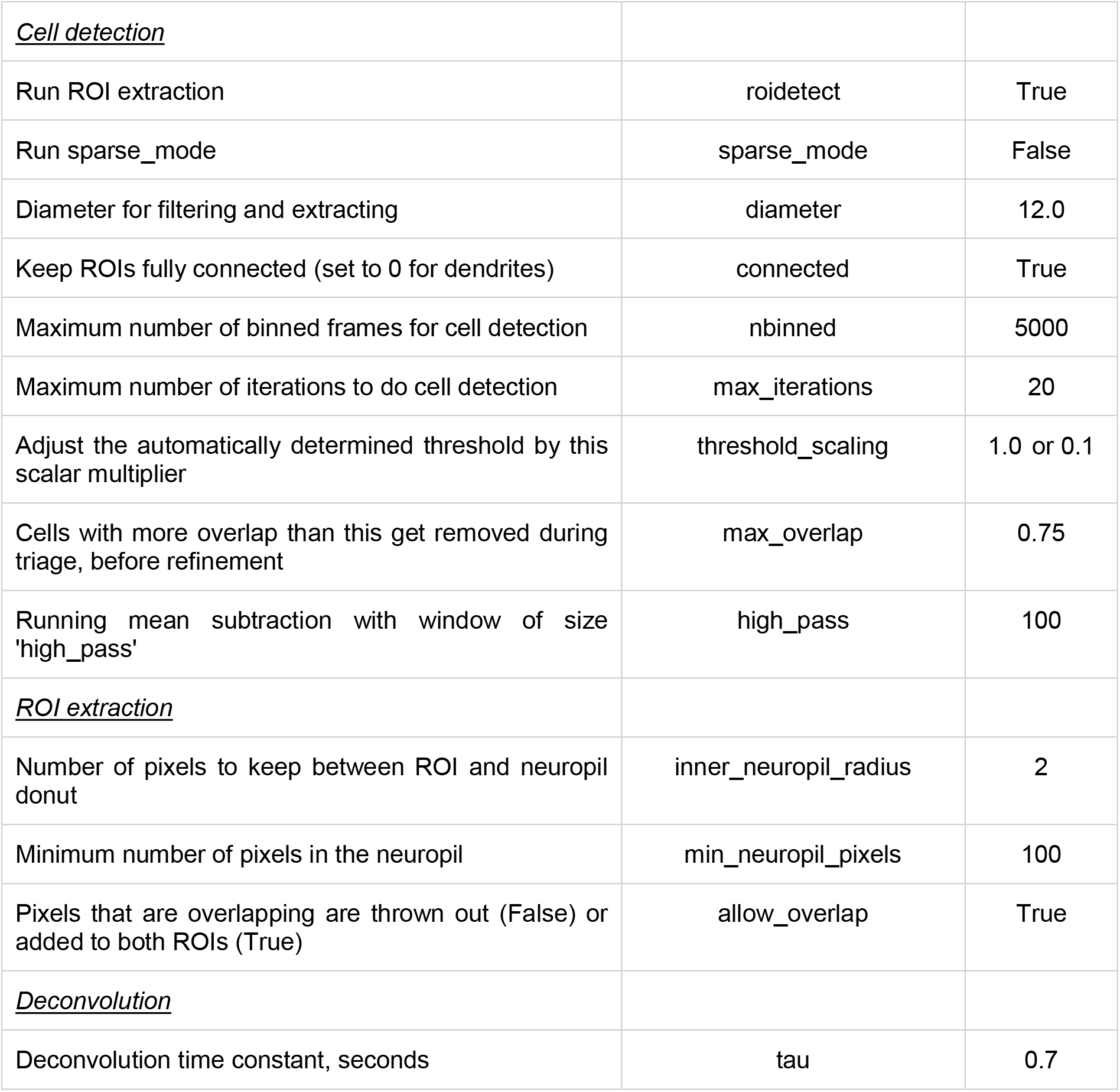

| Parameter | Variable | Value |
| --- | --- | --- |
| Sampling rate, frames per second | fs | 30 |
| <u>Registration</u> |  |  |
| Subsampled frames for finding reference image | nimg_init | 2000 |
| Number of frames per batch | batch_size | 200 |
| Maximum allowed registration shift, as a fraction of frame max(width and height) | maxregshift | 0.1 |
| Precision of subpixel registration (1/subpixel steps) | subpixel | 10 |
| Smoothing | smooth_sigma | 1.15 |
| Bad frames to be excluded | th_badframes | 100.0 |
| <u>Non-rigid registration</u> |  |  |
| Use nonrigid registration | nonrigid | True |
| Block size to register (** keep this a multiple of 2 **) | block_size | [128,128] |
| if any nonrigid block is below this threshold, it gets smoothed until above this threshold. 1.0 results in no smoothing | snr_thresh | 2.0 |
| maximum pixel shift allowed for nonrigid, relative to rigid | maxregshiftNR | 10 |
| <u>Cell detection</u> |  |  |
| Run ROI extraction | roidetect | True |
| Run sparse_mode | sparse_mode | False |
| Diameter for filtering and extracting | diameter | 12.0 |
| Keep ROIs fully connected (set to 0 for dendrites) | connected | True |
| Maximum number of binned frames for cell detection | nbinned | 5000 |
| Maximum number of iterations to do cell detection | max_iterations | 20 |
| Adjust the automatically determined threshold by this scalar multiplier | threshold_scaling | 1.0 or 0.1 |
| Cells with more overlap than this get removed during triage, before refinement | max_overlap | 0.75 |
| Running mean subtraction with window of size 'high_pass' | high_pass | 100 |
| <u>ROI extraction</u> |  |  |
| Number of pixels to keep between ROI and neuropil donut | inner_neuropil_radius | 2 |
| Minimum number of pixels in the neuropil | min_neuropil_pixels | 100 |
| Pixels that are overlapping are thrown out (False) or added to both ROIs (True) | allow_overlap | True |
| <u>Deconvolution</u> |  |  |
| Deconvolution time constant, seconds | tau | 0.7 |

The same analytical pipeline was applied to both the raw fluorescence traces as well as the deconvolved (“spikes”) signal, as extracted by the Suite2p toolbox. Generally, the raw fluorescence signal was preferred over the deconvolved one given that its extraction is more straightforward and relies on less assumptions. However, while the reported effects varied in magnitude depending on which of the two signals was considered, the same results were obtained on both datasets. The effects were entirely consistent. For raw signal analysis of each neuron, previous to any further step, we subtracted 0.7 of the corresponding neuropil fluorescence trace.

The number and height of calcium transients properties were calculated with the scipy function *find_peaks* on the raw calcium traces with the following parameters: height = 200, distance = 10 and prominence = 200. The decay was computed on the 10 best-isolated transients of each neuron, using the *OASIS* toolbox (https://github.com/j-friedrich/OASIS). We used the *deconvolve* function with the following parameters: penalty = 0, optimize_g = 10. Traces with an estimated decay over 2 seconds were considered cases of failed extraction and removed from further analysis.

The choice of the parameter values for transient detection is somewhat arbitrary. Similarly, it is debatable whether and how the calcium traces should best be normalized. Therefore, we tested the robustness of our findings by systematically varying signal extraction choices. We first varied the height and prominence threshold across a wide range of values (50 to 700 arbitrary units). We further computed transients features on normalized ΔF/F calcium traces. To normalize calcium signals, we used the baseline value as extracted by the *deconvolve* function. Also, in this case, we varied the height and prominence threshold across a wide range of values (0.5 to 3 arbitrary units). Finally, we computed two measures of neuronal activity that are independent of calcium transients detection: the average of the trace integral and its standard deviation, with and without normalization. Across all of these scenarios, the reported effects were robustly consistent.

Correlations were computed both as Pearson (numpy function *corrcoeff*) and Spearman (custom written function) coefficient on the z-scored signal. To both sets of coefficients, the Fisher correction (the inverse of the hyperbolic tangent function, numpy function *arctanh*) was applied. For power analysis, we first created a population activity vector by summing all the single neuron z-scored signals, and then estimated the power spectral density by applying the Welch method (sampling frequency = 30 Hz, number of points for fast Fourier transformation = 1024, no overlap, window length = 1 s).

For analysis of recovery from anesthesia, all recordings of the imaging session for a given animal were concatenated in Suite2p. As a consequence, each recording in the imaging session has the same set of reconstructed neurons. A time window of 5000 frames was used for the analysis to ensure continuous motionless periods. To track the neuronal activity changes, the number of fluorescence peaks, their amplitude, and the characteristic decay constant of the transients were considered. Each imaging session’s threshold was chosen to match the median activity in the pre-anesthesia (awake) state across all animals. To assess the relative changes of these parameters induced by anesthesia and their subsequent recovery over time, the parameters were normalized to their median value at the pre-anesthesia (awake) state. Notably, we focused our analysis on neurons that maintained some detectable activity during anesthesia, and neurons with no detected peaks were excluded from the distributions. Additionally, we applied the cut *decay constant* > 1/30 [s] (where 30 frames per second is an acquisition rate) to remove the traces where the OASIS algorithm considered a single noise peak to be a calcium transient.

Complexity analysis was performed in the Matlab (MathWorks) environment. For complexity analysis, we limited the number of neurons to the minimum (N_min_) present in any recording of any condition for each single mouse (median = 265, min = 156, max = 1068). The resulting matrix therefore had the T_rec_xN_min_ dimensions, where T_rec_ represents the time vector for the recording, with a length of 5 min and a sampling rate of 30 Hz. For recordings that had a number of neurons larger than N_min_ for that mouse, we randomly sampled n = N_min_ neurons and repeated the analysis 5 times. For every extracted parameter, we then considered the median value over the 5 repetitions. For further analysis, the signal was down sampled from the original sampling frequency of 30 Hz to 10 Hz (100 ms bins). The same analytical pipeline was then applied to both the raw fluorescence traces, as well as the deconvolved signal.

*tSNE clustering*. tSNE clustering was performed similar to [26]. Briefly, in a range between 5 and 45, the perplexity value that minimized the reconstruction error was selected. The number of PCA components used for this step was limited to 30. For the raw fluorescence signal, Euclidian distance was used, whereas for the deconvolved signal we opted for cosine distance, as it is better suited to a sparse signal. We computed the probability distribution of the resulting embedded matrix (2xT_rec_), that was then convolved with a 2D Gaussian window (standard deviation was set to be equal to 1/40 of the total maximum value). To evaluate the number of clusters in the distribution, we then applied a series of standard steps in image analysis: background subtraction with the rolling ball method, smoothing with a median filter, thresholding, watershedding to avoid undersegmentation, and extended minima transformation. Finally, the exterior boundaries of the objects were traced and counted. This gave the number of clusters.

##### Affinity Propagation Clustering (APC)

Affinity Propagation clustering was performed using a Matlab toolbox [https://www.psi.toronto.edu/index.php?q=affinity%20propagation] and similarly to [26]. We first obtained a distance map, which was computed as 1 minus the pairwise cosine distance between observations of the T_rec_xN_min_ matrix. This distance matrix was then fed to the affinity propagation algorithm with the input preference set equal to the median of the distance matrix.

##### Principal Component Analysis (PCA) clustering and variance explained

Standard PCA was applied to the T_rec_xN_min_ matrix. The number of clusters was computed as the number of components that was needed to cumulatively explain 90% of the variance of the input matrix. Further, we computed the loglog decay coefficient of number of components versus variance explained.

##### Community detection

To detect communities, we used the Louvain algorithm from the Brain Connectivity Toolbox (https://sites.google.com/site/bctnet/), a modularity maximization procedure widely used in studies examining brain networks [107]. This approach aims at subdividing the network into partitions that are more internally dense than would be expected by chance [53]. As input to the algorithm, we used Fisher-transformed correlation matrices obtained from calcium imaging time-series. Matrices were not thresholded, and both positive and negative correlations were taken into account to determine optimal modular partitions. The algorithm was evaluated while varying the resolution parameter gamma between 0 and 3, in steps of 0.1. For the multiresolution approach and hierarchical consensus clustering, data was analyzed using code available at https://github.com/LJeub/HierarchicalConsensus and according to the procedure described in [55]. The number of communities detected by the finest level partition of the consensus hierarchy was used for further analysis. While neurons in the awake condition tended to be spatially closer to each other than for the other conditions (Fig. S8E), this is unlikely to have influenced the results of the analysis, as the difference was minimal and there was no correlation between median distance in a recording and the number of detected communities (Fig. S8F).

#### Sleep scoring

Sleep scoring was carried out in two steps. We first used electrophysiological features (see below) to classify the behavioral state of the electrophysiological recordings. Then, using this dataset as ground truth, we extracted pupil/eyelid features that we used to extend our classification to the calcium imaging recordings.

##### Electrophysiological recordings

We divided the signal in 30s epochs with a 50% overlap, and used a rule-based approach similar to that applied in [57, 108]. NREM sleep epochs were defined as epochs having LFP power in the delta band (1-4 Hz) higher than the 70^th^ percentile, EMG broadband (30-300 Hz) power lower than the median, and no movement. REM sleep epochs were defined as epochs having a ratio between theta (6-12 Hz) and delta LFP power higher than the 70^th^ percentile, EMG broadband power lower than the 25^th^ percentile, and no movement. Finally, wakefulness epochs were defined as epochs having EMG broadband power that exceeded the 80^th^ percentile or with mouse movement. Given that this rule-based approach left ∼49% of the epochs as unclassified, we extended this classification with a machine-learning approach using the scikit-learn toolbox [109]. Using the classified epochs as the labelled dataset, we trained a K-nearest neighbors classifier (*number of neighbors=20, weights=distance, algorithm=auto, leaf size=5, p=2, scoring=f1 macro*) to which we fed the following quantile-transformed (*quantiles=20*) features: LFP power in the delta and theta band, ratio between LFP delta and theta power, EMG broadband power and average movement. After training, the algorithm was asked to predict the unclassified epochs. Predictions that were done with a probability estimate above 99% were kept, the others were left as unclassified. This adjunction to the rule-based approach allowed us to lower the percentage of unclassified (uncertain) epochs to ∼28%.

##### Pupil and eyelid analysis

During electrophysiological recordings, the eye of the mouse was recorded with a monochrome, infrared sensitive camera (UI-3360CP-NIR-GL Rev. 2, iDS imaging, Germany, objective: LMZ45T3 2/3” 18-108mm/F2.5 manual macro zoom lens, Kowa, Germany) under red light. Videos were captured with the uEye Cockpit software (iDS imaging, Germany) with a framerate of 30 Hz. Pupil, EMG and electrophysiological recordings were synchronized with a customized light/digital pulse shutter. During calcium imaging recordings were done with a, infrared sensitive camera (DMK 33UX249; The Imaging Source, Germany) equipped with a macro objective (TMN 1.0/50; The Imaging Source, Germany) at a frame rate of 10 Hz. Contours of the mouse eye were tracked using the deep neural network-based software module DeepLabCut [110] and subsequently processed in MATLAB. We trained a neural network (residual neural network, 152 layers, 200,000 iterations) to detect the upper, lower, left, and right edges of the pupil and eyelid, respectively, in images down-sampled to 256 pixels on the shorter edge (n = 1038 frames for videos from electrophysiology, n = 2255 frames for videos from calcium imaging). Besides the position of each tracked point, DeepLabCut provides a value quantifying the certainty about the determined position (which is low in the case of occluded objects, e.g. the pupil during an eyeblink). Samples with a certainty < 0.5 were linearly interpolated from the last point before to the first point after the respective samples which had a certainty of > 0.5 (0.24/0.56 % of total pupil samples and 0.11/0.12 % of total eyelid samples acquired during electrophysiology and calcium imaging experiments, respectively). We then calculated the pupil diameter (as the maximum distance between two opposing points of the pupil) as well as its center of mass, and the opening of the eye (as the distance between the top and the bottom eyelid). Finally, blinks were removed from the eye-opening data by linearly interpolating regions which exceed the moving median minus three times moving median absolute deviation (sliding window = 30 s).

##### Calcium imaging recordings

Using the expanded classification of the electrophysiology dataset, we extracted the following pupil/eyelid features: maximum and minimum pupil diameter, standard deviation of the pupil diameter, pupil area, pupil motion and eyelid distance. We then tested the extent to which it was possible to correctly predict the behavioral state on these features alone, similarly to Yüzgec et al., 2018. To this aim, we quantile transformed these features (*quantiles=[50, 100, 500]*), and passed them to a K-nearest neighbors classifier with similar hyper-parameters as the previous one (*number of neighbors=[5, 10], weights=uniform, algorithm=auto, leaf size=1, p=2, scoring=f1 macro)*. Hyper-parameter tuning was done using GridSearchCV. We then iteratively (n=100) tested the prediction accuracy on 25% of the dataset, yielding good average accuracy for wakefulness (∼86%) and NREM sleep (∼90%). On the contrary, most REM sleep was classified as NREM (∼62%) and the accuracy for this category was significantly lower (∼27%). Finally, we retrained the classifier on the entire dataset of pupil/eyelid electrophysiological features, and used it to predict the behavioral state of the calcium imaging dataset.

#### Two-Photon Spine Image Processing

In each animal, at least one GFP-expressing CA1 pyramidal neuron was selected and 1-3 dendrites of 20–50 µm length of each of the following types were analyzed: basal dendrites, oblique dendrites emerging from the apical trunk and tuft dendrites. Motion artefacts were corrected with a custom-modified Lucas-Kanade-based alignment algorithm written in Matlab. Spines that laterally emanated from the dendrite were counted by manually scrolling through the z-stacks of subsequent imaging time points of the same dendritic element, by an expert examiner blinded to the experimental condition. Protrusions from the dendrite that reached a threshold of 0.2 µm were scored as dendritic spines regardless of shape. If spine neck positions differed 0.5 µm on the subsequent images, the spine was scored as a new spine. Spines were scored as lost if they fell below the threshold of 0.2 µm. Spine density was calculated as the number of spines per µm. The turnover ratio was calculated for every time point by dividing the sum of gained and lost spines by the number of present spines. The survival fraction of spines was calculated as the percentage of remaining spines compared with the first imaging time point.

#### Statistical analysis

Statistical analyses were performed using R Statistical Software (Foundation for Statistical Computing, Vienna, Austria) or GraphPad Prism. All R scripts and datasets are available on GitHub https://github.com/mchini/Calcium-Imaging---Anesthesia. Nested data were analyzed with linear mixed-effect models to account for the commonly ignored increased false positive rate inherent in nested design [111]. We used “mouse”, “recording”, “neuron” (calcium imaging), and “single unit” (electrophysiology) as random effects, according to the specific experimental design. Parameter estimation was done using the lmer function implemented in the *lme4* R package [112]. Model selection was performed according to experimental design. Significance and summary tables for lmer model fits were evaluated with the *lmerTest* R package [113], using the Satterthwaite’s degrees of freedom method. Post hoc analysis with Tukey multiple comparison correction was carried out using the *emmeans* R package.

## Data Availability

Further information and requests for resources and reagents should be directed to and will be fulfilled by the corresponding author, J. Simon Wiegert.

### Data and Code Availability

The code generated during this study is available at https://github.com/OpatzLab/HanganuOpatzToolbox and https://github.com/mchini/Calcium-Imaging---Anesthesia

The calcium imaging and electrophysiology data sets generated during this study are available at https://gin.g-node.org/SW_lab/Anesthesia_CA1

