## Supplemental figures for "Anesthetics fragment hippocampal network activity, alter spine dynamics and affect memory consolidation"

**Fig. S1 – related to Fig. 1**

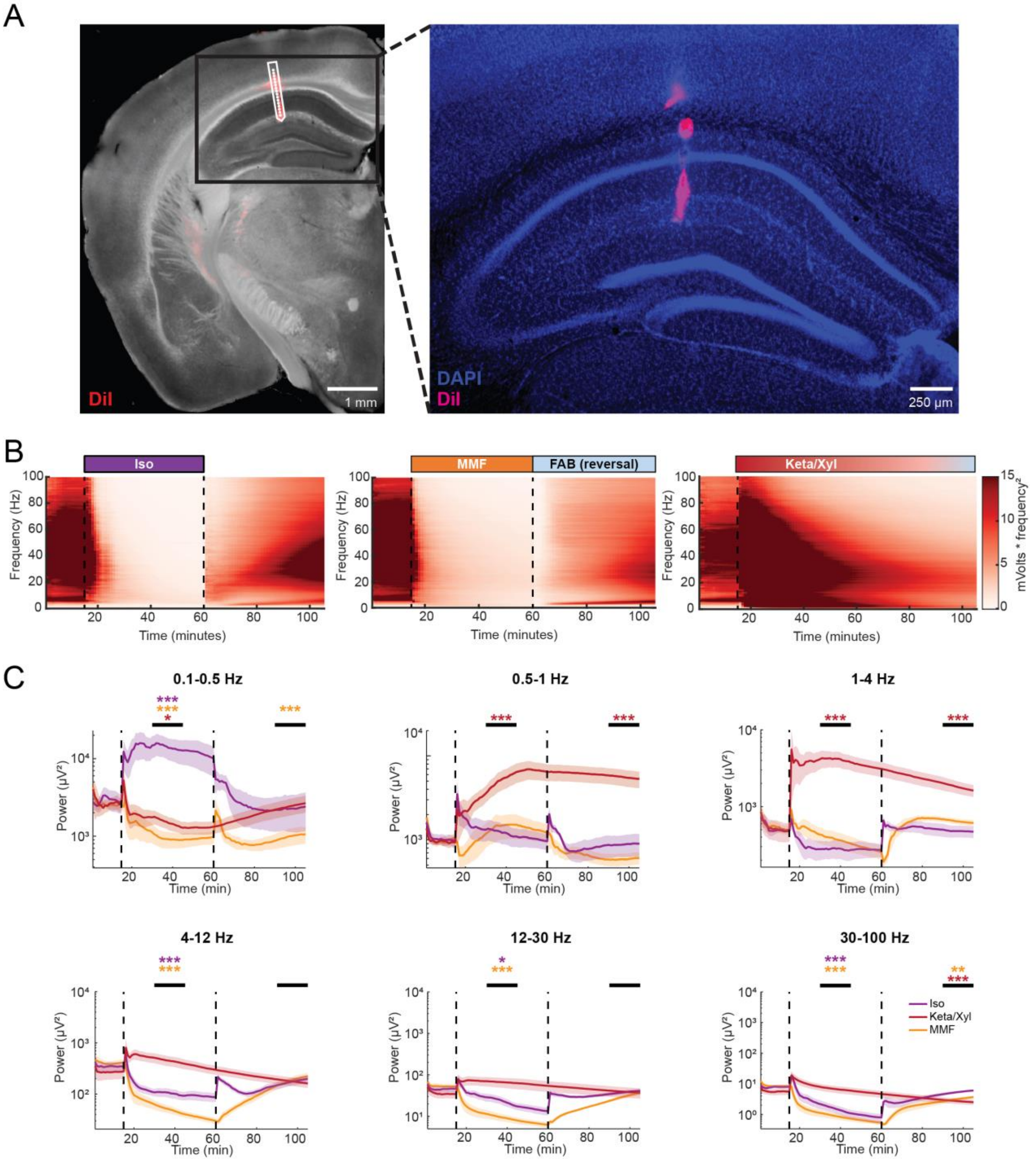


**Figure S1: LFP recordings in dorsal CA1 during wakefulness and anesthesia. (A)** Histology images showing the position of the recording electrode in the dorsal hippocampus. Left: Bright-field image merged with fluorescence image of DiI staining. The position of the electrode is indicated by white line drawing. Right: Close-up of dorsal hippocampus showing nuclear DAPI-staining in blue and DiI label from recording electrode position in magenta. **(B)** Heat map displaying raw LFP power in the 0-100 Hz frequency band for the three different anesthetic strategies. **(C)** Average LFP power over time in different frequency bands. Vertical dashed lines in all panels indicate time points of anesthesia induction (Iso, MMF, Keta/Xyl) and reversal (Iso & MMF only). Lines display mean ± SEM. Asterisks indicate significance of time periods indicated by black horizontal line compared to 15-min period before anesthesia. Anesthetic conditions are color-coded. * $p < 0.05$, ** $p < 0.01$, *** $p < 0.001$. For full report of statistics, see statistics table.

**Fig. S2 – related to Fig. 2**

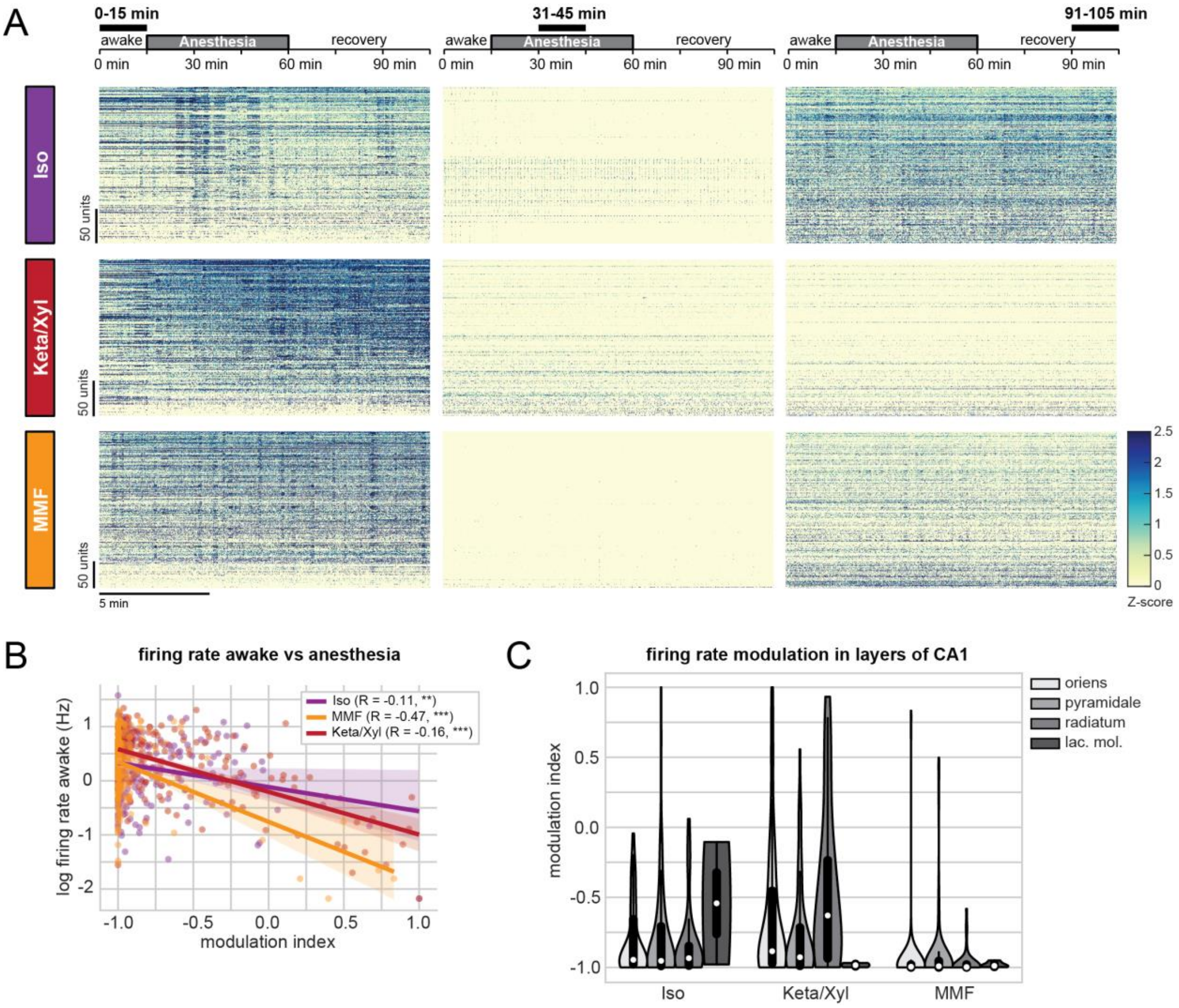


**Figure S2: Unit activity in dorsal CA1 during wakefulness and anesthesia. (A)** Raster plots of z-scored single-unit activity (SUA) during indicated time periods (black horizontal bars) for the three different anesthetic strategies. Units are sorted according to initial activity during wakefulness. Same data as in Fig. 2A. **(B)** Scatter plot showing modulation of unit activity during anesthesia with respect to firing rate during wakefulness. *** $p < 0.001$. **(C)** Violin plot showing modulation of unit activity during anesthesia with respect to CA1 anatomical layers. For full report of statistics, see statistics table.

**Fig. S3 – related to Fig. 3**

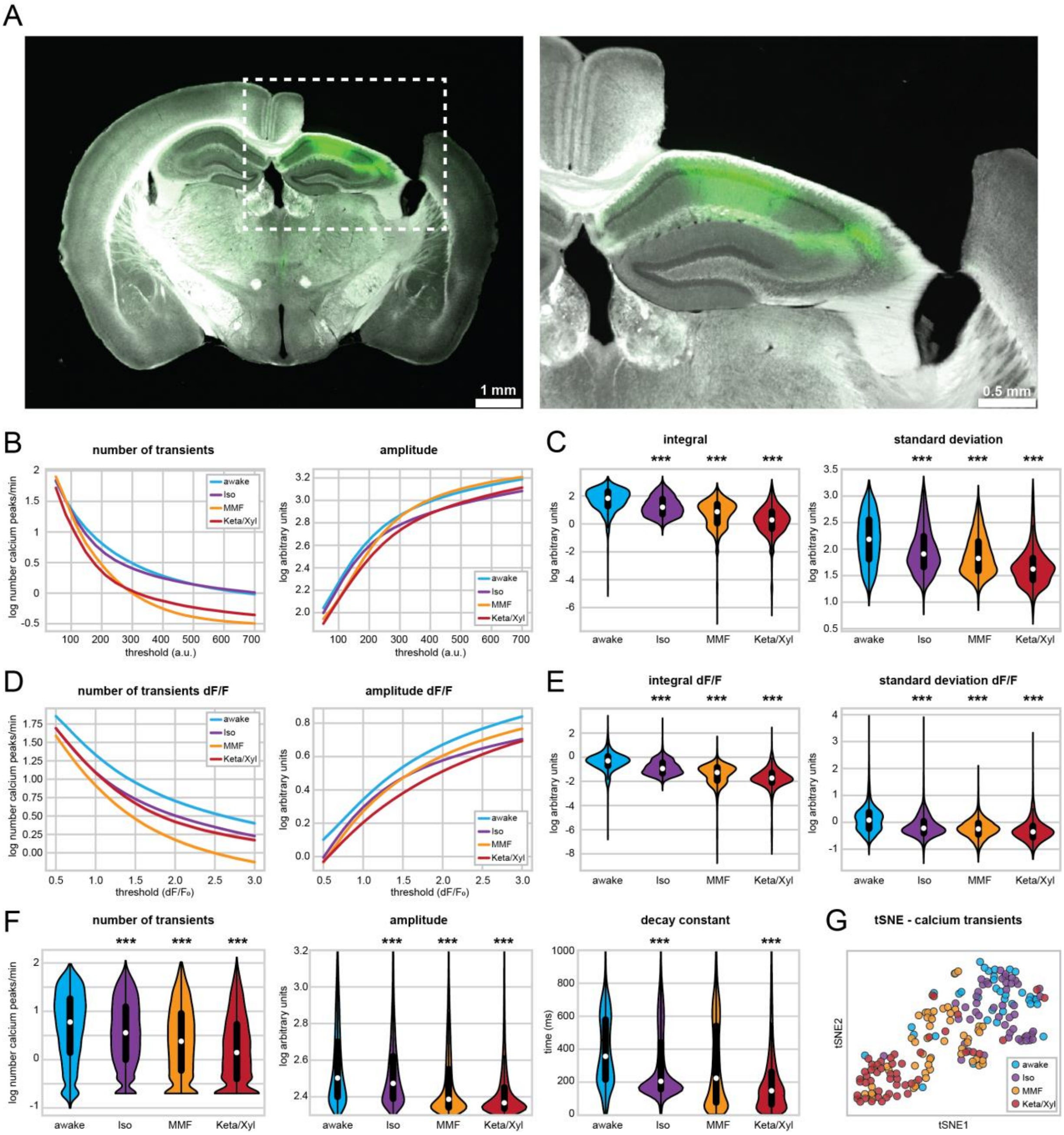


**Figure S3: Anesthesia-induced changes in calcium activity profiles are insensitive to the choice of signal extraction parameters. (A)** Histology images showing the position of the imaging window above the dorsal hippocampus. Left: Bright-field image merged with fluorescence. GCaMP6f expression is shown in green. Right: magnified view of the hippocampus. **(B)** Line plot of the number (left) and amplitude (right) of detected calcium transients across varying threshold values used for transient detection. **(C)** Violin plots quantifying the integral (left) and standard deviation (right) of calcium traces. White dots indicate median, vertical thick and thin lines indicate 1$^{st}$-3$^{rd}$ quartile and interquartile range, respectively. **(D, E)** As (B, C), for dF/F calcium transients and traces. **(F)** Violin plots quantifying the number (left), amplitude (middle), and decay (right) of calcium transients extracted with the *threshold_scaling* variable set to 0.1 instead of 1. **(G)** tSNE plot summarizing the average calcium transients properties. Each data point represents one recording session. Asterisks in (C, E, F) indicate significant differences to wakefulness. *** $p < 0.001$. Note, to facilitate readability, only differences to wakefulness are indicated. For full report of statistics, see statistics table.

**Fig. S4 – related to Fig. 3**

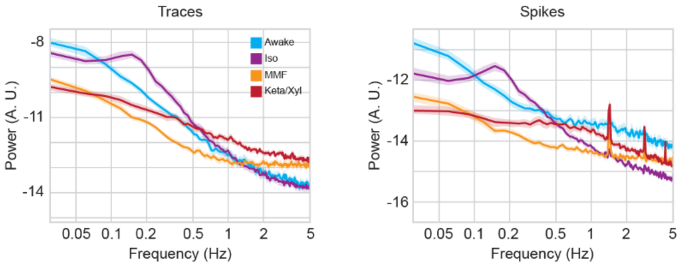


**Fig. S4. Oscillations of calcium transients are distinctly altered by Iso, MMF and Keta/Xyl.** Line plot displaying the spectrograms for population activity power, for raw calcium transients (left) and deconvolved spikes (right) during wakefulness and three different anesthetic conditions.

Fig. S5 – related to Fig. 4

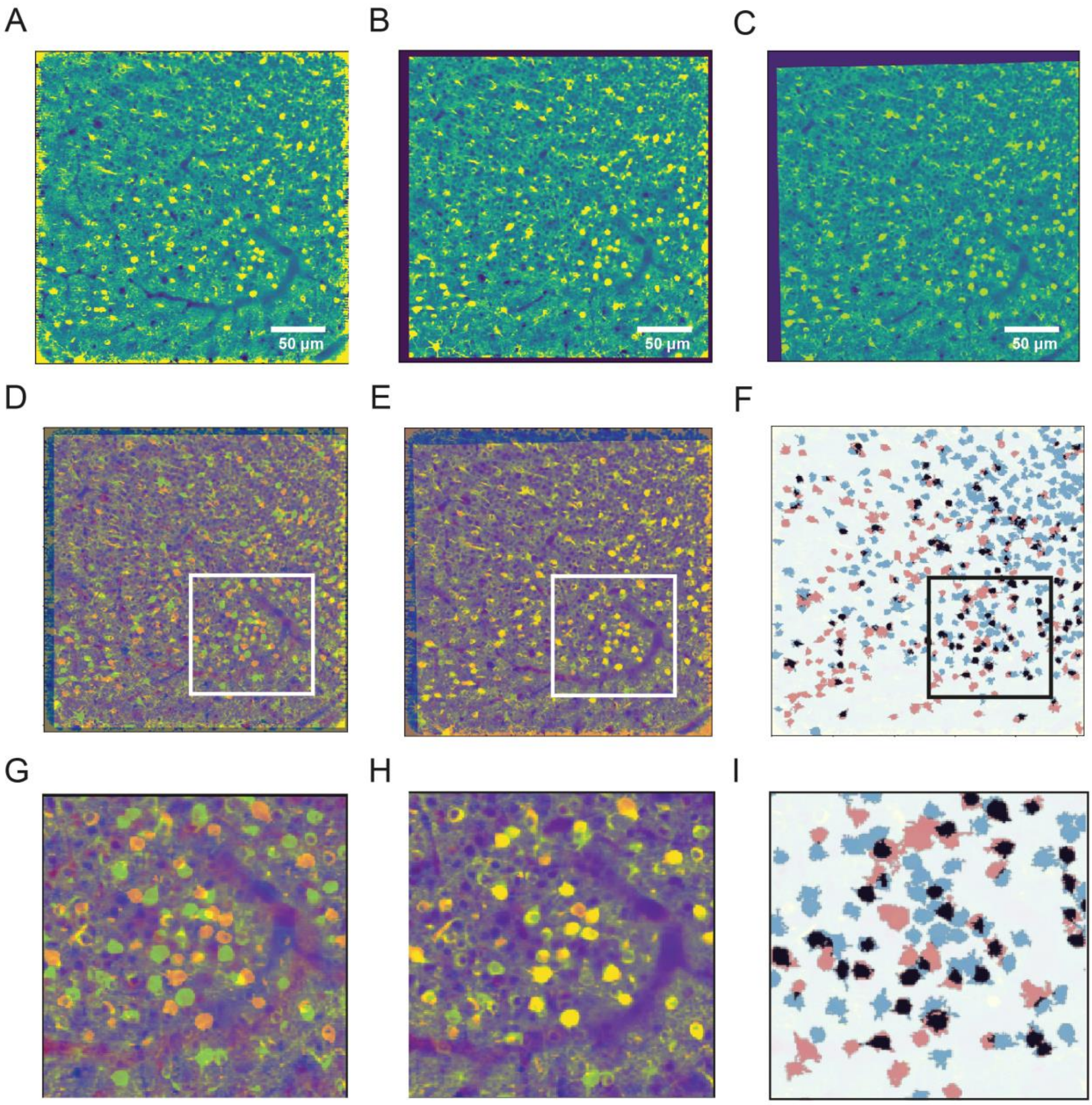


**Fig. S5: Alignment and ROI matching of imaging sessions.** Enhanced mean-intensity images are shown to demonstrate the alignment procedure between a recording in Iso **(A)** and one during wakefulness **(B)**. **(C)** Mean-intensity image of the awake recording aligned to the anesthesia condition. **(D)** Overlaid, enhanced mean-intensity images before and after **(E)** the alignment algorithm was applied. **(F)** ROIs of active neurons during wakefulness (pink), isoflurane anesthesia (cyan) and their overlap (black). **(G-I)** Magnified views of a randomly selected sections from panels (D) – (F).

### Fig. S6 – related to Fig. 4

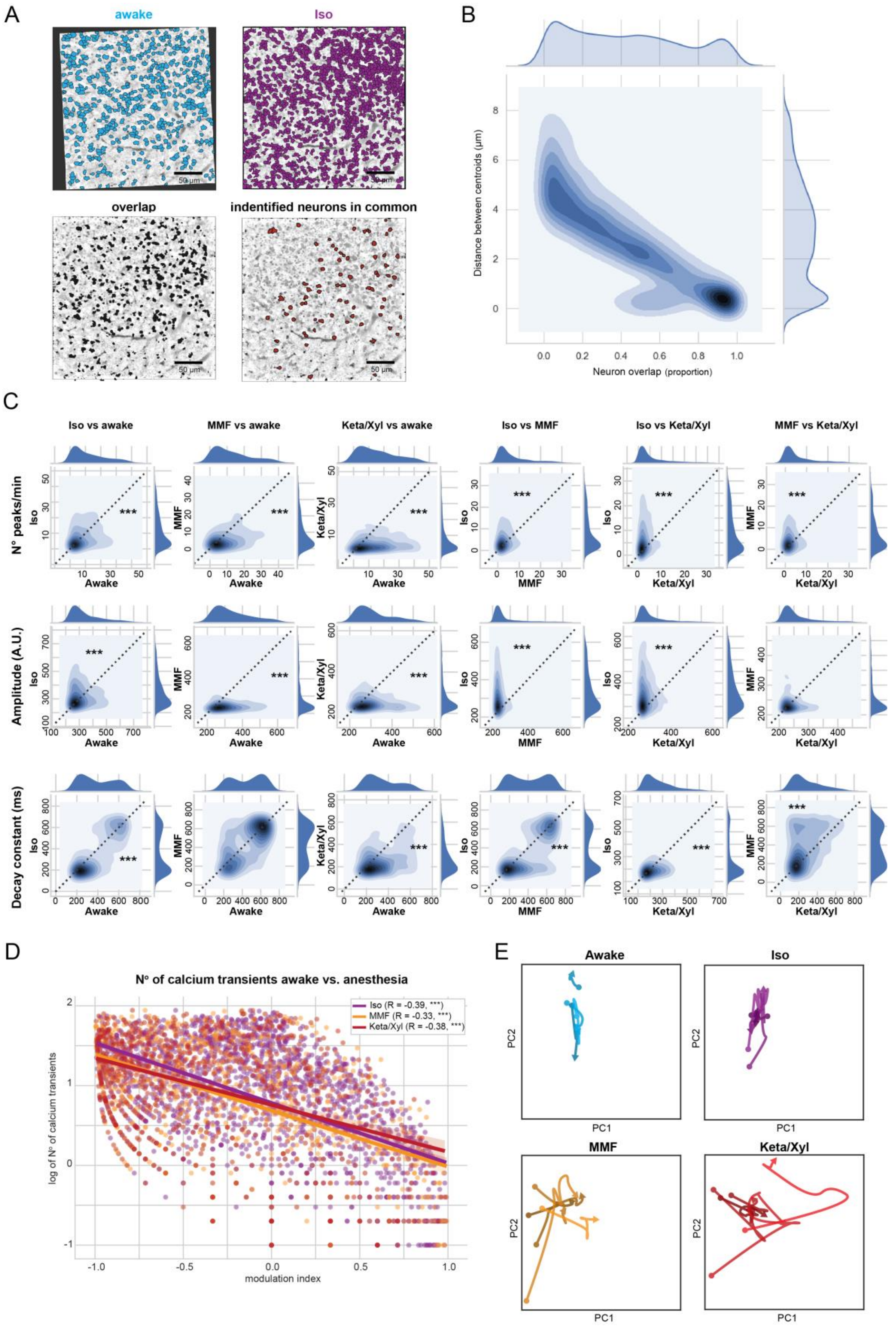


**Fig. S6: ID assignment and comparison of calcium transients in identified neurons between pairs of conditions. (A)** Time-averaged two-photon images of the same FOV in CA1 from an 'awake' (top left) and an Iso (top right) imaging session aligned to the Iso condition (same images as in figs. 3&4). ROIs of active neurons were automatically extracted with a lower quality threshold, accepting more neurons per recording (see Methods). Lower left: geometrical overlay of the two images with overlapping ROIs. Lower right: ROIs of identified neurons active in both conditions. **(B)** Kernel density plot and probability density functions for distances between centroids and area overlap for pairs of closest ROIs from the two different recordings. A clear bimodal distribution in both parameters is appreciable. Values in the lower right corner indicate highly matched neurons that were considered active in both conditions. **(C)** Kernel density plot and probability density functions for the number (top row), amplitude (middle row), and decay (bottom row) of detected calcium transients for each pair of conditions. From left to right: Iso vs awake, MMF vs. awake, Keta/Xyl vs. awake, Iso vs. MMF, Iso vs. Keta/Xyl, MMF vs. Keta/Xyl. Asterisks indicate significant differences between pairs of conditions. Asterisks are always on the side of the unity line with higher values. *** $p < 0.001$. **(D)** Scatter plot showing modulation of calcium transients during anesthesia with respect to calcium activity during wakefulness. *** $p < 0.001$. **(E)** Recovery of CA1 pyramidal neurons from anesthesia. The dots at the beginning of each trajectory represent median values of two principal components (PC1, PC2) calculated on the number, height and decay of the calcium transients during anesthesia (see Fig. 4E). The arrowheads represent the state 6 hours later. Blue dots correspond to awake states in the control group. Every trajectory connects anesthesia and post-anesthesia (recovery) states during the following 6 hours for a given animal. The trajectories were obtained using 3rd-degree B-splines.

**Fig. S7 – related to Fig. 5**

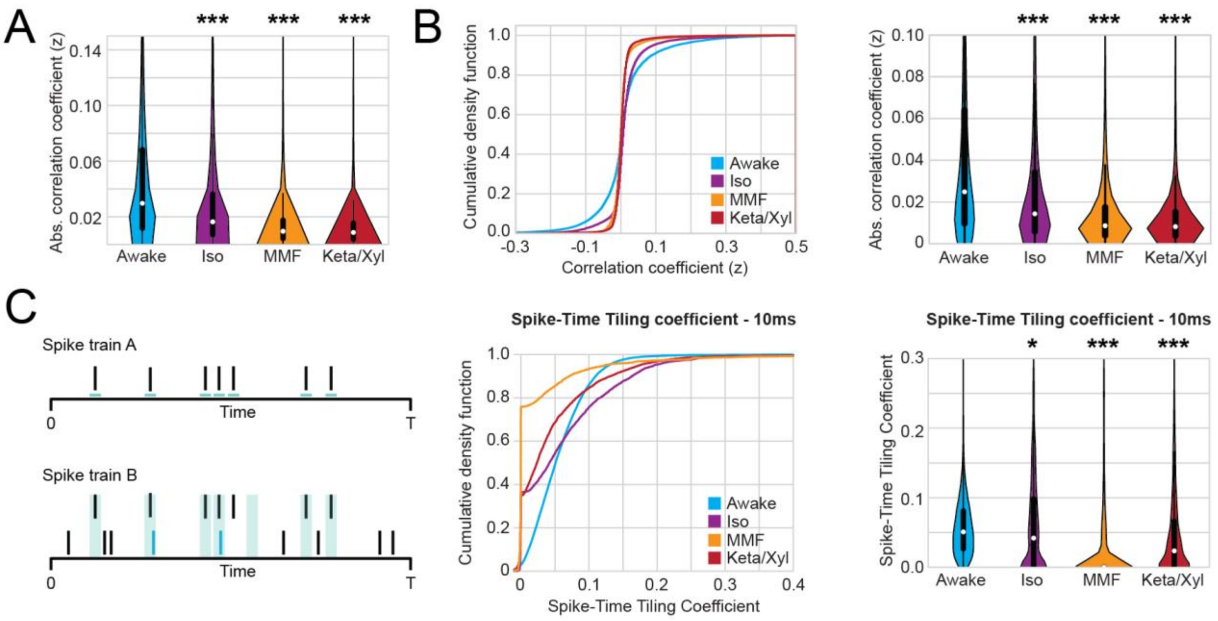


**Fig. S7. Additional correlation analysis. (A)** Violin plots quantifying absolute pairwise correlation of all neurons recorded during calcium imaging shown in Fig. 5. **(B)** Decorrelation of calcium activity during anesthesia is sustained also in neurons active during all conditions. Left: Line plot displaying cumulative distribution of Fisher-corrected Pearson correlation coefficients between the same pairs of neurons, which were active in each condition. Right: Violin plots quantifying absolute pairwise correlation coefficients. In violin plots, white dots indicate median, vertical thick and thin lines indicate $1^{st}$-$3^{rd}$ quartile and interquartile range, respectively. **(C)** Quantification of correlation between pairs of extracellularly recorded single units using the Spike-Time Tiling Coefficient with shorter integration window. Left: Schematic illustration the quantification of the Spike-Time Tiling Coefficient. Middle: cumulative distribution of the Spike-Time Tiling Coefficient with a 10 ms integration window. Right: violin plot quantifying the tiling coefficient. In all violin plots white dots indicate median, vertical thick and thin lines indicate $1^{st}$-$3^{rd}$ quartile and interquartile range, respectively. Asterisks indicate significant differences to wakefulness. * $p < 0.05$, *** $p < 0.001$. Note, only differences to wakefulness are indicated. For comparison between conditions, see statistics table.

**Fig. S8 – related to Fig. 6**

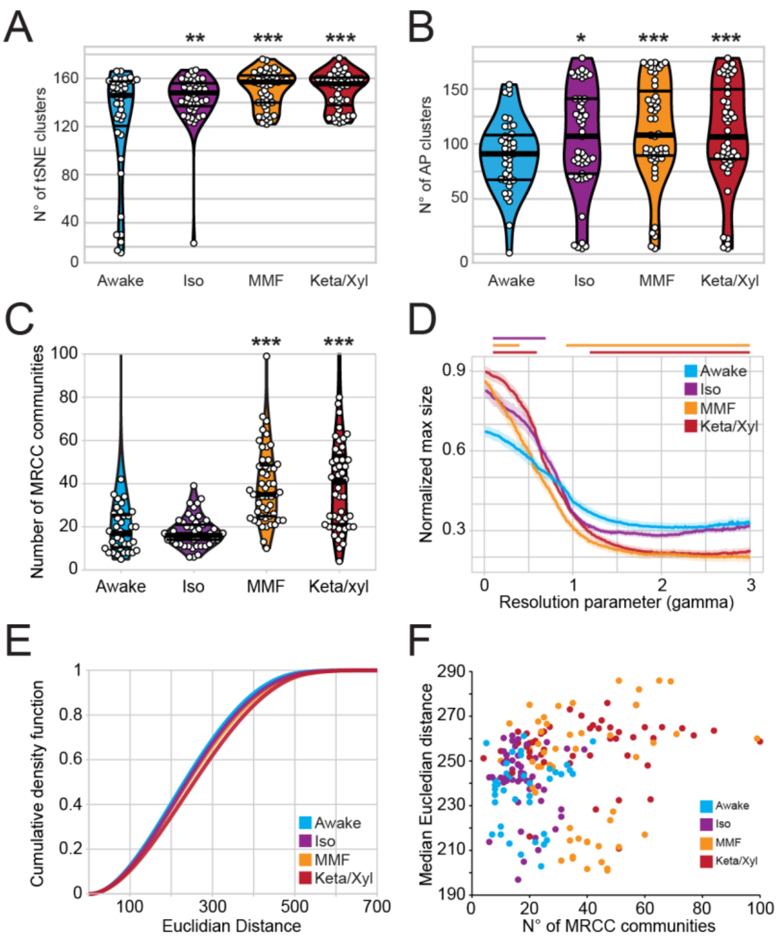


**Fig. S8. Clustering analysis of deconvolved calcium imaging data. (A)** Violin plot quantifying the number of tSNE clusters obtained from deconvolved calcium recordings (i.e., 'spikes') during wakefulness, Iso, MMF and Keta/Xyl anesthesia. **(B)** Violin plot quantifying the number of clusters obtained by affinity propagation from deconvolved calcium recordings (i.e. 'spikes') during the four different conditions. **(C)** Violin plot quantifying the number of communities obtained by multiresolution consensus clustering (MRCC) for the four different conditions. **(D)** Line plot quantifying maximum community cluster size normalized by the total number of neurons across the resolution parameter gamma ranging from 0 to 3. **(E)** Cumulative distribution of the distance between neurons randomly selected for the spatial cluster analysis showing no difference between conditions. **(F)** Scatter plot displaying absence of a relationship between the median distance of neurons and the number of detected communities. Horizontal lines in violin plots indicate median and 1$^{st}$-3$^{rd}$ quartile. Asterisks in (A) – (C) indicate significant differences to wakefulness. * $p < 0.05$, ** $p < 0.01$, *** $p < 0.001$. Horizontal lines above plots in (D) - (E) indicate significant difference to wakefulness. Anesthetic conditions are color-coded. Note, only differences to wakefulness are indicated. For comparison between conditions, see statistics table.

Fig. S9 – related to Fig. 7

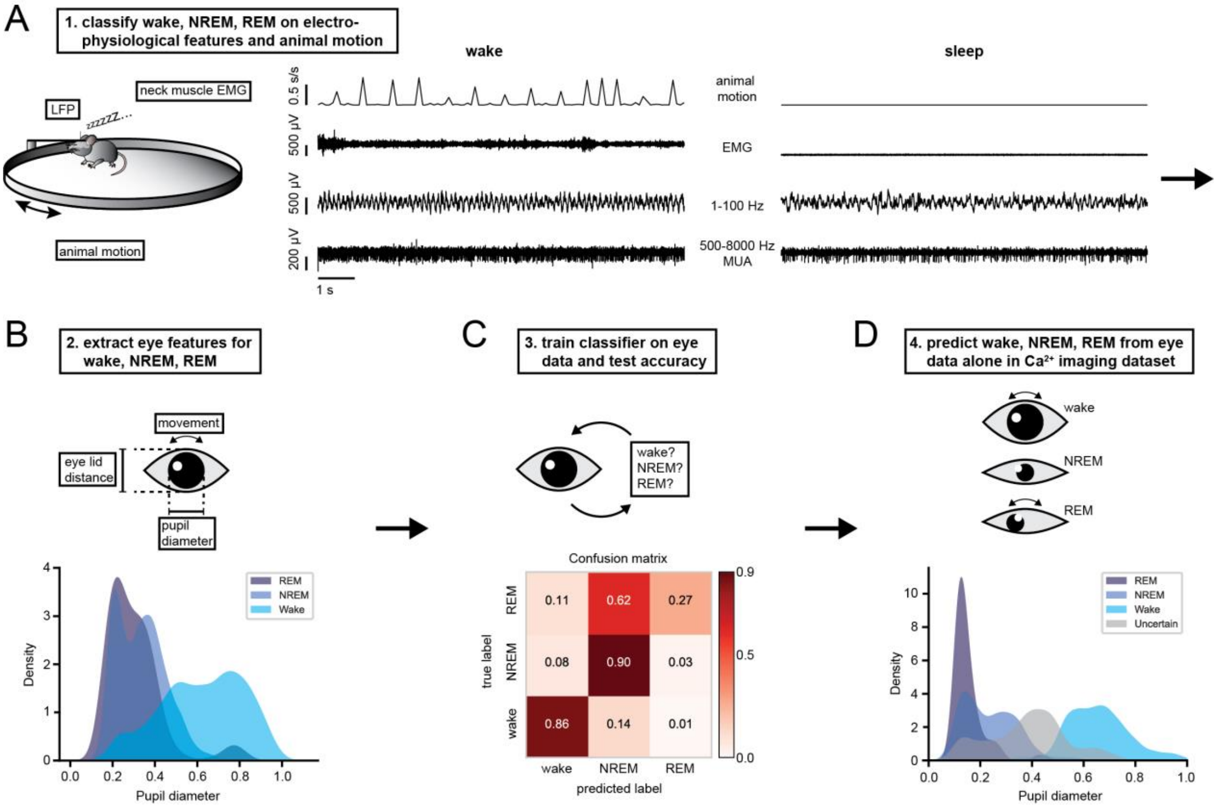


**Fig. S9. Sleep scoring.** Sleep scoring was carried out in two steps. We first used electrophysiological features to classify the behavioral state of the electrophysiological recordings. Then, using this dataset as ground truth, we extracted pupil/eyelid features that we used to expand our classification to the calcium imaging recordings. **(A)** Wake, NREM and REM sleep epochs were classified based on animal motion, neck muscle EMG broad power, MUA and several LFP features. **(B)** Using the electrophysiology-based classification, the following pupil/eyelid features were extracted to classify sleep: maximum and minimum pupil diameter, standard deviation of pupil diameter, pupil area, pupil motion and eyelid distance. **(C)** Confusion matrix and prediction accuracy for the classification of sleep periods based on eye imaging alone. **(D)** Probability distribution of pupil diameter for predicted wake, NREM and REM periods in the calcium imaging dataset.

**Fig. S10 – related to Fig. 8**

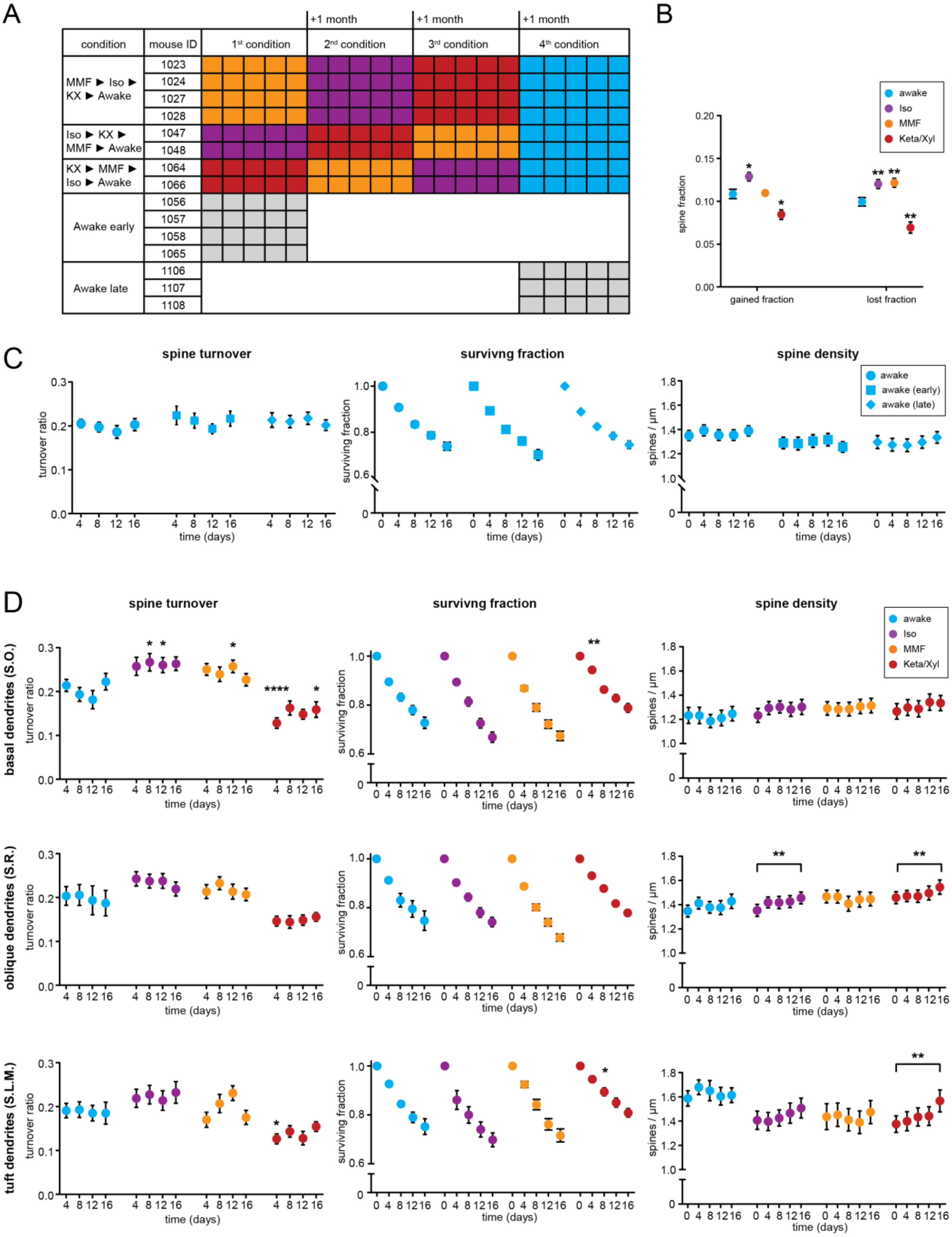


**Fig. S10. Chronic spine imaging. (A)** Experimental scheme for chronic spine turnover measurements. Spine imaging was performed in a pseudo-randomized order for the different anesthetics followed by imaging during wakefulness. Each colored box indicates one imaging session. For each condition, imaging was done five times every four days, followed by a one-month break. To control for long-term effects of anesthesia and age on the awake condition, we performed imaging only during wakefulness in additional mice as indicated. **(B)** Dot plots showing quantification of overall gain and loss of spines during chronic imaging under the four different treatments. Dots indicate mean ± SEM. **(C)** Dot plots showing quantification of spine turnover (left), spine survival (middle) and spine density (right) during wakefulness after anesthetic treatments (same data as "awake" in Fig. 7B) and wakeful imaging in the two control groups ("awake early", "awake late"). Dots indicate mean ± SEM. **(D)** Dot plots showing quantification of spine turnover (left column), spine survival (middle column) and spine density (right column) separately for basal (top row), oblique (middle row) and tuft dendrites (bottom row). Same data as in fig. 8B. Asterisks indicate significant differences to wakefulness. * p < 0.05, ** p < 0.01.

**Fig. S11 – related to Fig. 9**

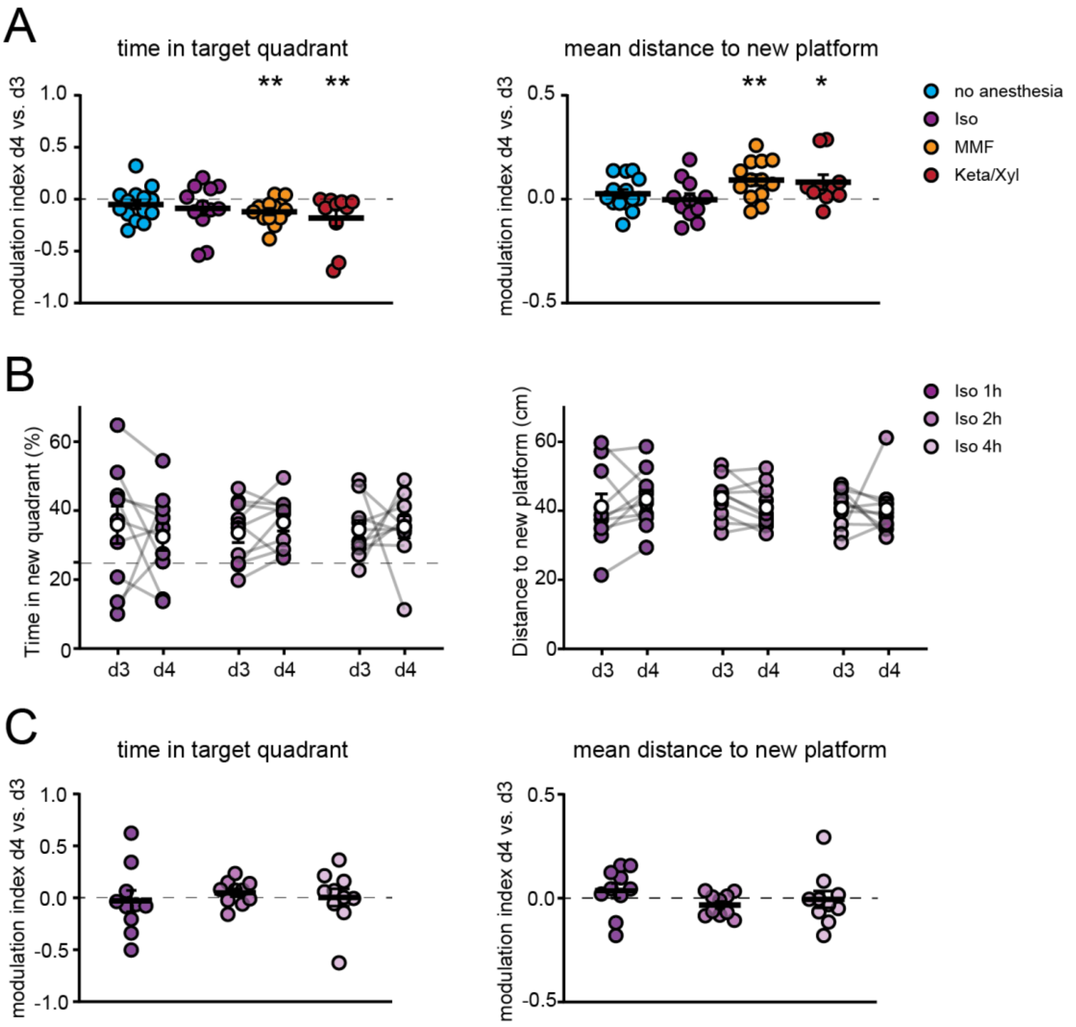


**Fig. S11. Comparison of the effect of Iso, MMF and Keta/Xyl on episodic memory consolidation. (A)** Scatter plots showing quantification of change in the time spent in the new target quadrant (left) and distance to the new platform (right) on day 4 after 1 h of indicated anesthesia or no treatment compared to day 3. **(B)** Scatter plots showing quantification of time spent in the new target quadrant (left) and distance to the new platform (right) after reversal learning on day 3 and on day 4 after 1, 2, and 4 h of Iso anesthesia. Filled, colored circles indicate individual animals, White circles indicate mean ± SEM. **(C)** Scatter plots showing quantification of change in the time spent in the new target quadrant (left) and distance to the new platform (right) on day 4 after 1, 2, and 4 h of Iso anesthesia compared to day 3. Asterisks indicate significant deviation from 0. * p < 0.05, ** p < 0.01. Note, significant differences between groups were not evident.
